# MultiomicsTracks96: A high throughput PIXUL-Matrix-based toolbox to profile frozen and FFPE tissues multiomes

**DOI:** 10.1101/2023.03.16.533031

**Authors:** Daniel Mar, Ilona M. Babenko, Ran Zhang, William Stafford Noble, Oleg Denisenko, Tomas Vaisar, Karol Bomsztyk

## Abstract

**Background:** The multiome is an integrated assembly of distinct classes of molecules and molecular properties, or “omes,” measured in the same biospecimen. Freezing and formalin-fixed paraffin-embedding (FFPE) are two common ways to store tissues, and these practices have generated vast biospecimen repositories. However, these biospecimens have been underutilized for multi-omic analysis due to the low throughput of current analytical technologies that impede large-scale studies.

**Methods:** Tissue sampling, preparation, and downstream analysis were integrated into a 96-well format multi-omics workflow, MultiomicsTracks96. Frozen mouse organs were sampled using the CryoGrid system, and matched FFPE samples were processed using a microtome. The 96-well format sonicator, PIXUL, was adapted to extract DNA, RNA, chromatin, and protein from tissues. The 96-well format analytical platform, Matrix, was used for chromatin immunoprecipitation (ChIP), methylated DNA immunoprecipitation (MeDIP), methylated RNA immunoprecipitation (MeRIP), and RNA reverse transcription (RT) assays followed by qPCR and sequencing. LC-MS/MS was used for protein analysis. The Segway genome segmentation algorithm was used to identify functional genomic regions, and linear regressors based on the multi-omics data were trained to predict protein expression.

**Results:** MultiomicsTracks96 was used to generate 8-dimensional datasets including RNA-seq measurements of mRNA expression; MeRIP-seq measurements of m6A and m5C; ChIP-seq measurements of H3K27Ac, H3K4m3, and Pol II; MeDIP-seq measurements of 5mC; and LC-MS/MS measurements of proteins. We observed high correlation between data from matched frozen and FFPE organs. The Segway genome segmentation algorithm applied to epigenomic profiles (ChIP-seq: H3K27Ac, H3K4m3, Pol II; MeDIP-seq: 5mC) was able to recapitulate and predict organ-specific super-enhancers in both FFPE and frozen samples. Linear regression analysis showed that proteomic expression profiles can be more accurately predicted by the full suite of multi-omics data, compared to using epigenomic, transcriptomic, or epitranscriptomic measurements individually.

**Conclusions:** The MultiomicsTracks96 workflow is well suited for high dimensional multi-omics studies – for instance, multiorgan animal models of disease, drug toxicities, environmental exposure, and aging as well as large-scale clinical investigations involving the use of biospecimens from existing tissue repositories.

## BACKGROUND

Multi-omics is a relatively new discipline that aims to reverse engineer biological systems by a) acquiring large molecular datasets for different ome layers of intracellular organization (1) and b) computationally integrating these heterogeneous datasets to gain a deeper understanding of phenotypes (2). The exponential growth of the multi-omics field has been fueled, in part, by advances in high throughout analytical technologies such as next generation sequencing, mass spectrometry, and others (1,3,4). Over the last decade the exponential increase in multiome publications has outpaced each one of the single-omic studies including genomics, epigenetics, transcriptomics, epitranscriptomics, proteomics, and metabolomics (5).

Historically, methods for effective profiling of tissues’ molecular complexity in health and disease have not until now been readily available. This past technological inadequacy is being mitigated by improvements in multi-omics analytical and computational tools, allowing multiome analysis of the same tissue samples (4,6-8). Integration of multidimensional datasets with phenotypes offers new opportunities to better understand intracellular regulation of organs’ physiology and dysfunction (3,9) as well as organism development (9), growth, and aging (10,11). Still, until now, multiome profiling has consisted of no more than three omic datasets from a single biospecimen (1,12), yielding an incomplete view of intracellular information flow and limiting the ability to define molecular causality.

Freezing is the preferred way to store and transport biospecimens for research (13-15). In clinical settings, formalin fixation and paraffin embedding (FFPE) is the most commonly used method for longitudinal tissue specimen storage. FFPE blocks are also commonly used to transport clinical samples for molecular diagnosis (e.g. MGMT methylation in glioblastoma (16)). All clinical laboratories in the US are required to hold onto diagnostic tissue blocks (mostly FFPEs) for several years, greatly multiplying the number of human tissue specimens in archives. The estimated hundreds of millions of FFPE tissue samples acquired over the years provide a vast resource for the discovery of disease molecular pathways linked to histopathology. Frozen and FFPE tissue blocks have been used in single-omic studies including genomic (17), transcriptomic (18,19), proteomic (20,21), and epigenomic (22-25) studies. Still, such biospecimens remain highly underutilized in the multi-omics field (23,26) because sample and analyte preparation methods are inefficient, slow, labor intensive, and low throughput (23), let alone the challenge of using a single biospecimen to generate multidimensional profiles. Thus, better tools and methods are needed to retrieve chromatin, DNA, RNA, and protein from biospecimens such as frozen and FFPE tissues to advance omics research in health and disease.

We have previously developed a 96-well microplate sonicator, PIXUL, that offers unmatched sample preparation throughput capabilities for a broad range of high throughput analytical workflow applications (27-29). Here, we took advantage of mouse organs’ epigenomic, transcriptomic, epitranscriptomic, and proteomic differences as substrates to develop systems that integrate 96-well plate format sample preparation, PIXUL, and analytical, Matrix (30-32), platforms for multiome profiling of FFPE and frozen tissues, MultiomicsTracks96.

## MATERIALS, DEVICES AND METHODS

Hardware/labware (**Table S1**) and kits/enzymes (**Table S2**) catalog numbers and commercial suppliers are listed in the supplementary tables.

### Reagents

Isopropanol (3223-0010) from Acros Organics. Ethanol (2716) from Decon Labs. Trifluoroacetic acid (TFA) (A116-50) from Fisher Chemical. O.C.T. compound (4585), sodium chloride (NaCl) (S-271-3) and Triton X-100 (BP151) from Fisher Scientific. Dulbecco’s Modified Eagle Medium (DMEM SH30021.0) from HyClone. Penicillin/streptomycin (P/S) (15749), phosphate buffered saline (PBS) (70013-032), TRIzol (15596018), and UltraPure Distilled Water (10977-015) from Invitrogen/Life Technologies. Fetal bovine serum (FBS 43635-500) from Jr. Scientific. Acetonitrile (9017-03) and chloroform (9180-01) from JT Baker. NP40 (198596) from MP Biomedicals. cOmplete Tablets Mini (04693159001) from Roche. Dithiothreitol (DTT) (D0632), EDTA (E3134), Tris–HCl (T3253), IGEPAL CA-630 (I8896), sodium deoxycholate (30970), and tetramethylammonium bromide (TMAB) from Sigma.

### Buffers

Preparations of all buffers and stock solutions were done with nuclease free reagents and UltraPure distilled nuclease free H_2_O. PBS: 137 mM NaCl, 10 mM Sodium phosphate, 2.7 mM KCl, pH 7.4; TE: 10 mM Tris-HCl, 1 mM EDTA, pH 7.5; Immunoprecipitation (IP) buffer: 150 mM NaCl, 50 mM Tris–HCl (pH 7.5), 5 mM EDTA, NP-40 (0.5% vol/vol), Triton X-100 (1.0% vol/vol); Elution buffer-Proteinase K: 25mM Tris Base, 1% IP Buffer, 1mM EDTA, 80μg/ml Proteinase K; Proteinase K buffer (PK buffer): 10mM Tris-HCl pH 8.0, 10mM EDTA, 0.5% SDS, 500μg/ml Proteinase K, and 40mM DTT.

### Devices

#### CryoGrid system

The complexity and heterogeneity of disease pathways require analysis of large numbers of tissue samples. Almost any disease, and often therapeutic intervention, are systemic conditions where animal models provide the means to understanding multiorgan dysfunction (33). To store and sample large numbers of biospecimens for multi-omics analysis, we designed a platform for cryostoring multiple tissue samples and engineered a hand-held rotary tool for rapid sampling of frozen tissues, CryoGrid system, which consists of CryoBox, CryoBlock, thermometer/thermocouple, QR barcoded CryoTrays, and CryoCore (34).

#### PIXUL

A 96-well plate sample preparation sonicator for multi-omics applications (Matchstick Technologies, Inc, Kirkland, WA and Active Motif, Carlsbad, CA) (27).

### Methods

#### Frozen and FFPE mouse organs

Post-mortem brains, hearts, kidneys, livers, lungs, and muscle were used from 12-week-old C57bl/6 mice.

#### Tissue freezing, sampling, jetting cores into PIXUL plate, and sonicating

Before freezing tissues, a 24-well CryoTray was placed into the chilled CryoBlock maintained at < -70°C in a CryoBox filled with dry ice pellets. Fresh post-mortem tissues were immediately put on ice and then one by one were placed in individual pockets of the 24-well CryoTray with a small amount of embedding matrix (e.g., OCT or Leica CryoGel) injected into the bottom of the wells for rapid freezing and immobilizing of tissue fragments. The total amount of time to freeze 24 tissues was less than 20 minutes. Frozen tissues were placed in the wells and immobilized with either OCT or Leica CryoGel. The CryoTray, with tissues covered by a QR code labeled lid, was stored at -80°C (34).

#### Tissue formalin fixing and paraffin embedding (FFPE), microtome sectioning and extracting rehydrated samples for RNA and DNA based analysis

Fresh frozen mouse organs were fixed with 10% formalin overnight and then embedded in paraffin. Following fixation, the tissue samples were processed through dehydration, clearing, and infiltration. Dehydration was initiated in 70% ethanol, progressing through two changes of 95% ethanol and then three changes of 100% ethanol. The tissues were then cleared by three changes of xylene and infiltrated by melted paraffin wax. FFPEs were sectioned using a microtome, and slices were deparaffinized and rehydrated using serial ethanol dilutions.

#### Chromatin isolation from frozen tissues

Before sampling, the CryoTray with frozen tissues was inserted in a chilled CryoBlock (< -70°C) in a CryoBox filled with dry ice pellets. An iPad was used to display a Google Sheet document with the organ layout legend and convenient access to metadata to facilitate sampling with the CryoCore. Before coring, the CryoCore trephine was cooled by plunging it into the dry ice pellets. One to two CryoCore cores extracted from frozen tissue were jetted with PBS (drawn from the CryoCore syringe reservoir) directly into 1.5ml tubes on ice. After PBS was aspirated, 1% Formaldehyde (2.5% Formalin) was added to the tube and tissue fragments were cross-linked for 30min at room temperature. Tubes were centrifuged at 6500xG for 2min (RT), supernatants were discarded, 0.5ml 125mM Glycine was added to stop the cross-linking, and tubes were vortexed and incubated for 5min (RT). After centrifugation at 6500xG (2min, RT), PBS was added to wash the pellets, and the tubes were centrifuged at 10,000xG (2min, RT). Then, supernatants were discarded and the pellets were either frozen and cryostored (-80°C) or resuspended in 100μl chromatin shearing buffer on ice. Samples were transferred to 96-well round-bottom plate (Corning Costar 3799) and were sheared in PIXUL (Pulse=50, PRF=1.0, Burst=20, T=4min x 4). Sheared chromatin samples were transferred to 0.5ml tubes on ice and centrifuged at 16,000xG for 15min (4°C). The supernatants contained soluble chromatin suitable for immediate use in Matrix ChIP (31,32), and any unused portions were aliquoted (to minimize thawing and freezing) and stored at -80°C for later use.

#### Chromatin isolation from FFPE tissue blocks

10-20 5μm thin slices were cut with microtome and collected in 1.5mL tubes. Paraffin was removed using SafeClear (1.2mL) incubating at 50°C for 3min, and then tubes were centrifuged at 10,000xG for 2min. Samples were rehydrated with serial dilutions of 100%, 50%, and 10% ethanol, 10min each (30 min total time). After each incubation, tubes were centrifuged at 10,000xG for 2min. Pellets were resuspended in 200μL extraction buffer, incubated at 95°C (20min), allowed to cool briefly, and centrifuged, and 1mM PMSF and 0.5mM DTT (final concentrations) were added to each tube. Aliquots of 100μL each were transferred to 96-well PIXUL plate and sonicated (Pulse=50, PRF=1.0, Burst=20, T=4 min x4), then transferred to tubes, centrifuged 16,000xG for 15min (4°C), and stored at -80°C.

#### DNA isolation from frozen tissues

One CryoCore core extracted from frozen tissue was jetted with PBS (drawn from the CryoCore syringe reservoir) directly into 1.5ml tubes on ice. PBS was removed and 100μL of 25mM Tris Base pH 10.2 + 1uL IP buffer + 3uL RNase (10mg/mL stock) was added to the tubes, which were incubated for 30min at RT. 4μL Proteinase K (20mg/mL stock) was added to each tube, and contents were transferred to wells of 96-well plate and processed in PIXUL (no pre-chilling, Pulse=50, PRF=1.0, Burst=20, T=30min). Plates were briefly centrifuged, and contents were transferred to 1.5mL tubes which were then boiled for 10min, chilled on ice, and centrifuged at 16,000xG for 15min. The supernatants containing ssDNA were used immediately in MeDIP or stored frozen (-20°C) for future use.

#### DNA isolation from FFPE tissue blocks

10-20 5μm thin slices were cut with microtome and collected in 1.5mL tubes. Paraffin was removed using SafeClear (1.2mL) incubating at 50°C for 3min, and then tubes were centrifuged at 10,000xG for 2min. Samples were rehydrated with serial dilutions of 100%, 50%, and 10% ethanol, 10min each (30 min total time). After each incubation, tubes were centrifuged at 10,000xG for 2min. Pellets were resuspended in 200μL extraction buffer, incubated at 95°C (20min), allowed to cool briefly, and centrifuged. After 300ng/μL RNase was added to each tube, samples were incubated for 30min at RT, and then 800ng/μL Proteinase K was added. The suspensions (100μl aliquots/well) were transferred to PIXUL plate, sonicated (Pulse=50, PRF=1.0, Burst=20, T=30min), transferred to 0.5mL tubes, boiled for 10min, chilled on ice, and centrifuged at 16,000xG for 15min (4°C). The supernatants containing ssDNA were used immediately in Matrix methylated DNA immunoprecipitation MeDIP (28,31,35) or stored frozen (-20°C) for future use.

#### RNA isolation from frozen tissues

One CryoCore core from frozen tissue was jetted with PBS (drawn from the CryoCore syringe reservoir) directly into 1.5ml tubes on ice. PBS was removed, 100μL TRIzol was added to the tubes, and samples were transferred to 96-well PIXUL plate and then sonicated (Pulse=50, PRF=1.0, Burst=20, T=6min). Well contents were transferred to 1.5ml tubes and additional TRIzol was added to 1mL total volume. RNA was isolated using standard TRIzol protocol (ThermoFisher), and dry pellets were resuspended in 50μl UltraPure water and stored at -80°C.

#### RNA isolation from FFPE tissue blocks

10-20 5μm thin slices were cut with microtome and collected in 1.5mL tubes. Paraffin was removed using SafeClear (1.2mL) at RT (10min), and then tubes were centrifuged at 10,000xG for 2min. Samples were rehydrated with serial dilutions of 100%, 50%, and 10% ethanol, 10min each (30 min total time). After each incubation, tubes were centrifuged at 10,000xG for 2min. Pellets were resuspended in 200μl extraction buffer. Samples were divided into 100μl aliquots in 1.5ml tubes, incubated at 95°C for 20min, and allowed to cool briefly. After centrifugation, 1.6μg/μL Proteinase K was added, and samples were transferred to 96-well plates and sonicated in PIXUL (Pulse=50, PRF=1.0, Burst=20, T=6min). Well contents were transferred to 1.5mL tubes to proceed with TRIzol RNA isolation as above.

#### DNase I Treatment (for RNA isolation)

After the TRIzol step, MgCl_2_ and CaCl_2_ (2.5mM each) were added to each 50μL sample. Next, DNase I was added (0.1U/μL), and samples were incubated at 37°C for 30 minutes. Finally, 5mM EDTA was added to each sample, and then samples were incubated at 75°C for 10 minutes to inactivate DNase I. Finally, 5mM MgCl_2_ was added in order to chelate the EDTA.

#### 96-well plate matrix quantitative reverse transcription real time PCR (Matrix RT-qPCR)

Isolated RNA (100ng) was reverse transcribed with Superscript IV, 0.2 mM dNTP (GeneScript, 95040-880), and random hexamers in 10μl reactions in 96-well microplates for 10 minutes at 50°C then 10 minutes at 80°C. RT reactions were diluted 10-fold with elution buffer prior to running qPCR. Housekeeping genes were used to normalize qPCR results (36). RT-qPCR primers are listed in supplementary **Table S3**. In-house software, PCRCrunch, was used to acquire, store and analyze qPCR data sets generated by Matrix RT-qPCR (30).

#### mRNA-sequencing (RNA-seq)

After isolation, RNA was run through Zymo RNA Clean & Concentrator. Sequencing libraries were prepared using Zymo-Seq RiboFree Total RNA Library Kit with RNA between 205-480ng and libraries amplified between 13-14 cycles of PCR as per manufacturer’s protocol. Library validation was performed as described below. Libraries were diluted as per Illumina protocol to a final pooled loading concentration of 650pM in resuspension buffer (RSB) plus Tween 20 with a 10% PhiX spike-in and sequenced in Illumina P2 cartridges on NextSeq 2000 that employed a dual-index, paired-end, 61 base read length (PE61).

After sequencing, Fastq files were downloaded from Illumina Basespace. Pre-processing was performed using fastp, and the quality of the runs was validated using fastqc (37). The pre-processed Fastq files were then aligned to the mouse mm10 genome and converted to BAM files using HISAT2 (paired-end library, reverse correspondence of read to transcript) (38).

#### 96-well plate Matrix ChIP-qPCR/seq

Microplate-based Matrix chromatin immunoprecipitation (ChIP)-qPCR was done as previously described where ChIP antibodies were attached to wall wells via Protein A (28,32). To minimize the background, wells were blocked with 5% BSA and single stranded Salmon DNA. After chromatin immunocapture, ChIP cross-linking was reversed and DNA was eluted using Proteinase K at 55°C and then boiled, yielding ssDNA used in qPCR (27,32). Antibodies are listed in **Table S4.**

Modified protocol was used for Matrix-ChIP-seq where, after immunocapture, cross-linking was reversed overnight at 55°C in the presence of Proteinase K (400ng/µL) and DNA was extracted using phenol/chloroform, yielding dsDNA. Sequencing libraries were generated using Next Gen DNA Library Kit, Active Motif (Carlsbad, CA). Library validation was performed as described below. Libraries were sequenced using NextSeq 2000.

#### 96-well plate Matrix-MeDIP-seq

96-well microplate-based Matrix methylated DNA immunoprecipitation (Matrix-MeDIP) of ssDNA was done as previously described where 5mC antibody was attached to well walls via Protein A (28,31). Well walls were blocked with 5% BSA. PIXUL fragmented ssDNA was used as input. After washes, immunocaptured ssDNA was recovered by incubation with Proteinase K (200ng/µL) in 100µL/well of elution buffer (55°C for 45min, 95°C for 10min, then 4°C). MeDIP was validated with PCR. Sequencing libraries were prepared using xGen ssDNA library kit (IDT). Library validation was performed as described below. Libraries were sequenced using NextSeq 2000.

#### 96-well plate Matrix-MeRIP-qPCR/seq

96-well microplate-based Matrix methylated RNA immunoprecipitation (Matrix-MeRIP) of RNA was done using a protocol similar to Matrix-MeDIP where m6A or m5C antibody was attached to wall wells via Protein A. Blocking buffer containing RNaseOUT in a 1:100 ratio (v/v) was used for the antibody-RNA incubation step. Fragmented RNA was used as input, and immunocaptured RNA was recovered by incubation with Proteinase K (200ng/µL) in 100µL/well of elution buffer (55°C for 10min, 95°C for 10min, then 4°C). Samples were concentrated from 50μL to 16μL and DNase I treated (Zymo RNA Clean & Concentrator). RT-qPCR was used for QC. Zymo-seq RiboFree Total RNA Library Kit was used to generate dsDNA indexed libraries. Given the low amounts (<50ng) of RNA, the RiboFree Universal Depletion was carried out over 16hrs. Library Index PCR was done using 16-18cycles. Library validation was performed as described below. Libraries were sequenced using NextSeq 2000.

#### Library validation

After library generation for each method (RNA-Seq, Matrix ChIP-seq, Matrix MeDIP-seq, and Matrix MeRIP-seq), each library was validated, with Collibri Library Quantification Kit PCR used to verify adapter attachment, organ-specific-primer PCR used to show retained specificity, and Agilent 4200 TapeStation used to assess base pair length and library quality.

#### Protein isolation from frozen tissues

Fresh frozen samples were sampled using the CryoCore and resuspended in protein extraction buffer (buffer M3: 50mM 50 mM HEPES 8.5, 5% glycerol, 500mM NaCl, 1% IGEPAL CA-630, 2% sodium deoxycholate, 1% SDS, 1% Benzonase supplemented with complete protease inhibitors panel, Roche cOmplete) (39). After removal of all liquid, the tissues were resuspended in extraction buffer M3. Proteins were extracted using PIXUL (Pulse=50, PRF=1.0, Burst =20, Time=10min). Samples were centrifuged at 16,000xG for 15 min (4°C) and supernatant was collected. The protein concentration was measured using BCA assay.

#### Protein isolation from FFPE tissue blocks

Sixty 5μm microtome curls were cut from mouse organ FFPEs (2 blocks per tissue) and combined in 1.5mL tubes. The sections were twice deparaffinized using xylene (10 min at 37°C), and rehydrated by serial incubations in 100%, 70%, and 50% ethanol (5 min each). After removal of the last solution, the samples were incubated first in 0.1% formic acid (30 min) and washed in 100mM Tris pH 10. They were then incubated in minimal volume of 100mM Tris pH 10 (1h at 95° C with mixing at 600rpm), temperature was decreased to 60°C, and samples were incubated for another hour (mixing 600rpm) in Thermomixer (Thermo). After removal of all liquid, the tissues were resuspended in extraction buffer M3. Proteins were extracted using PIXUL (Pulse=50, PRF=1.0, Burst=20, Time=10min). Samples were centrifuged at 16,000xG for 15 min (4°C) and supernatant was collected. The protein concentration was measured using BCA assay.

#### Peptide digestion, mass spectrometry (MS) analysis, and data processing

Extracted proteins (10μg) were digested using an SP3 protocol on Sera-Mag SpeedBead Carboxylate-Modified beads (E3 and E7, 1:1)(40), using acetonitrile as the organic solvent. The proteins were digested in-situ with mixture of trypsin/Lys-C mix (Promega, WI) (1:10 w/w enzyme to proteins) for 16hrs at 37°C. The digest was cleaned up in-situ on the SP3 beads and eluted with 50μL of 0.5% DMSO, acidified with 5μL of 1% formic acid and dried down. For tandem mass tag (TMT) labeling, the samples were reconstituted in 10μL of 20% acetonitrile in 100mM tetramethylammonium bromide (TMAB) buffer for 1h vortexing. TMTPro reagents (ThermoFisher) were dissolved in 20μL of anhydrous acetonitrile. The 2μL of the reagents was added to each of the 16 samples, carefully mixed and incubated with mixing for 1hr at room temperature. To quench the reaction, 2μL of 2M Tris pH 8.0 was added and incubated for 45min. The TMT labeled sample was dried down, reconstituted in 1% acetonitrile, 0.1% TFA and fractionated using Pierce High pH Reversed-Phase Peptide Fractionation Kit. Fractions were dried down and stored at -20°C until analysis when they were reconstituted in 40μL of 5% acetonitrile, 0.1% formic acid.

#### LC-MS/MS analysis

After desalting on a C18 trapping column (Reprosil-Pur 120 C18-AQ, 5µm, 0.1 x 40mm, Dr. Maisch HPLC GmbH, Germany) (flow rate 4µL/min), the digested peptides were separated on an analytical column (Reprosil-Pur 120 C18-AQ, 5µm, 600×0.075mm, Dr. Maisch HPLC GmbH) maintained at 50°C. The following multi-step linear gradient was used: 2-9%B in 14min, 9-31%B in 151min, 31-44%B over 24min, and at the end of the gradient column was washed with a ramp to 80%B and re-equilibrated (A - 0.1% formic acid in water, B – 80% acetonitrile, 0.1% formic acid, flow rate of 0.4µL/min). An LC-MS/MS consisting of an EasyLC 1200 (Thermo Scientific, CA), and a Thermo Orbitrap Exploris 480 (Thermo Fisher, San Jose, CA) mass spectrometer with electrospray ionization was used for the analysis. Data-dependent acquisition consisting of one MS1 scan and a set of data-dependent MS2 scans with total cycle time 3sec was performed with following parameters: full MS1 scan 300-1800 m/z at 120,000 resolution, max ion time 50ms, normalized AGC 100%, and data-dependent MS2 scan with 30sec dynamic exclusion after one MS2 scan for a given precursor at 10ppm window. DDA MS2 was accomplished using TurboTMT mode of the Xcalibur v4.0 implemented on the Orbitrap Exploris 480 with isolation window 0.7amu and MS2 scan at 30,000 resolution, normalized HCD collision energy 32, max ion time 120ms, and normalized AGC target of 200%.

#### Data processing

The peptides and proteins were identified using TMT16 workflow in Fragpipe (v19.1) against UniprotKB release 2022_12 (22-Dec-2022) mouse reference proteome supplemented with common contaminants and 1:1 decoys (22086 protein sequences, 44172 total entries). Search parameters included mass accuracy 20ppm, trypsin KR/P specificity with maximum 2 missed cleavages, variable modifications: oxidation (M), fixed modification: carbamidomethyl (C) with FDR 0.01 for both PSM and protein identifications. TMT quantification data was further processes in R (v.4.2.1patched). The raw reporter ion intensities were normalized to the total reporter ion intensity across all proteins and log2 transformed. Missing values were replaced by 0. For correlation analysis, duplicate samples were averaged and correlations between FFPE and fresh frozen were calculated using Pearson method.

#### Segway genome segmentation

The Segway software (41) was applied to FFPE and frozen samples separately. In each sample prep, genomes across the four organs are concatenated and binned at 250bp resolution. Each analysis considered four epigenomic assay types (ChIP-seq of H3K27Ac, H3K4m3, and Pol II, as well as MeDIP-seq of 5mC). For computational efficiency, each model is trained on chromosome 1 to segment the genome into 15 labels, and then the trained model is applied to all chromosomes.

Super-enhancers were downloaded from corresponding organs in the dbSUPER databset, where cortex and cerebellum annotations are concatenated to yield brain-specific super-enhancer annotations (42). LiftOver was applied to convert mm9 super-enhancer annotations to mm10 genome coordinates (43). Segtools (44) was used to calculate the overlap of each genomic segmentation to super-enhancers. Among the 15 labels assigned by Segway, the label with the highest enrichment of super-enhancers is deemed as predicted super-enhancer annotations, and an F1 score is calculated based on the precision and recall of super-enhancer prediction of that label in each organ and sample prep.

To derive organ-specific super-enhancers that are captured by both FFPE and frozen samples, for each organ, we retain a list of predicted super-enhancer genomic regions that are not predicted in other organs, with separate lists for FFPE and frozen samples. Comparing these lists to the organ-specific superenhancers from dbSUPER yields lists of novel organ-specific super-enhancer predictions.

#### Protein expression prediction

Protein quantities were first normalized by library size and then subjected to log transformation. Other types of assays were summarized and mapped to genes, with read counts converted to log2-counts-per-million and normalized using the TMM (trimmed mean of M values) method (45). This procedure yielded 5,572 proteins, each with eight types of measurements. To predict protein quantities, we fit a linear regression model based on the other seven measurements, as well as Boolean indicators of sample prep (frozen vs. FFPE) and organ type. To evaluate the performance of the model, we used five-fold cross-validation, randomly splitting the proteins into five folds and iteratively training on four folds and testing on the held-out fold. Predictions for the five held-out folds were then concatenated together, and the Pearson correlation and mean squared error (MSE) between the predicted and original protein quants were calculated for each organ and sample prep. The overall Pearson correlation and MSE were also calculated across all organs and sample preps. To assess the predictive power of different types of multi-omics measurements, we also performed linear regression based on subsets of predictors. To compare the predictive power of different sets of predictors, we performed one-sided paired Wilcoxon signed-rank tests on the Pearson correlation coefficients and MSEs across cross-validation folds, organ, and sample prep. The resulting p values were subjected to Benjamini-Hochberg correction.

## RESULTS

### A high-throughput system for storing and sampling of frozen tissues (CryoGrid (34)) integrated with PIXUL-based RNA and chromatin analyses in 96-well plate-based assays (Matrix) (Fig.1)

PIXUL offers unparalleled sample preparation throughput to explore multiple samples in one experiment (27), but the upstream steps such as cryostorage and particularly tissue sampling represented a challenge in the workflow of multi-omics analysis. To mitigate this bottleneck, we designed a high-throughput platform for storing frozen tissues and sampling, CryoGrid, which includes the CryoCore (34). CryoCore is a hand-held rotary tool for rapid sampling of frozen tissues for molecular analysis and histology. With CryoCore, the same frozen tissue can be sampled multiple times, yielding reproducible core sizes (34). We used mouse brain, heart, kidney, liver, lung, and muscle fragments to integrate and optimize CryoGrid and CryoCore with PIXUL protocol to extract RNA and chromatin from frozen tissue samples (**Fig.1A**). One core (1-2mm) was used for each of the following: H&E histology **(Fig.1B),** RNA isolation followed by RT-qPCR, and ChIP (Matrix ChIP-qPCR). In Matrix-RT and Matrix-ChIP assays, we used primers to genes specific to brain (*Syn1*), heart (*Tnnt2*), kidney (*Fxyd2*), liver (*Alb*), lung (*Sftpa1*), and muscle (*Tnnc2*) to assess the quality of the methods **(Fig.1C).** Matrix ChIP was done using antibodies to RNA polymerase II (Pol II), histone H3, H3K27Ac, H3K4m3 and IgG (background control). In our previous work we have found that Pol II ChIP signal intensity at 5’ ends of genes recapitulates mRNA levels (27,30,32). Consistently, results of CryoGrid/CryoCore-based assays demonstrated excellent agreement between the levels of mRNAs, Pol II, and permissive H3K27Ac and H3K4m3 histone modifications assayed in these organs **(Fig.1D-E).** Further, these results demonstrate that a PIXUL-tissue-ChIP assay of just one epigenetic mark (either H3K27Ac or H3K4m3) measured at 5’ ends of six organ-specific genes is sufficient to identify the mouse organ (brain, heart, kidney, liver, lung, or muscle) without looking at the histology **(Fig.1B)** We used CryoGrid-PIXUL frozen organs data as a benchmark for PIXUL-FFPE method development (below).

**Fig.1.**
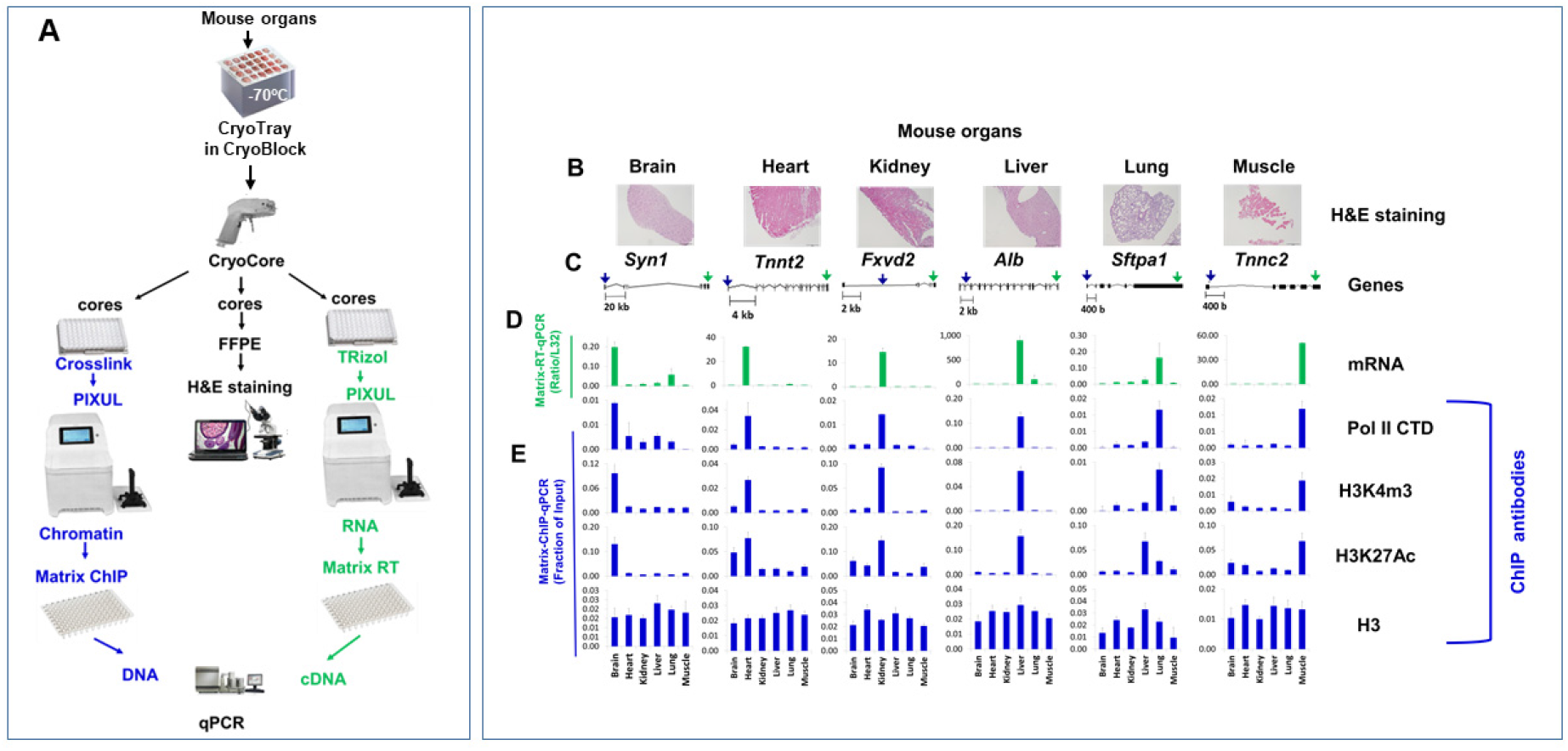
CryoGrid system integrated with PIXUL: A high-throughput system for rapid sampling of frozen tissue for histology, transcript, and epigenetic analysis. ***A.*** CryoTray with multiple frozen mouse organs in CryoBlock (<-70°C). Each row contains organs from the same mouse; from left to right, brain, heart, kidney, liver, lung, and muscle. A single tray holds all six organs from four mice. CryoCore is a battery-operated, easy-to-use tool to reproducibly extract tissue cores (1-2mm long and 1mm in diameter) from frozen tissues*. Left,* cores were jetted into wells of 96-well PIXUL plates, crosslinked, and sonicated in PIXUL. PlateHandle (black), next to PIXUL, is a hand-held tool for picking up plates, thus avoiding contamination and the need to handle plates directly with hands. Sheared chromatin was used in Matrix ChIP, *left*. *Center,* cores were used to prepare FFPE blocks for H&E staining. *Right,* shows the protocol for RNA extraction. ***B.*** CryoCore cores from CryoTray mouse frozen brain, heart, kidney, liver, lung, and muscle stained with H&E (Haemotoxylin and Eosin). ***C.*** Cartoons of the genes. Blue arrows show positions of qPCR primers used in ChIP and green arrows show the positions of qPCR primers used in RT-qPCR. ***D.*** CryoCore-PIXUL extracted RNA was assayed in Matrix RT-qPCR using primers to 3’ ends of the genes. ***E.*** Cores from the frozen organs were cross-linked and sonicated. PIXUL-sheared chromatin samples were analyzed for RNA Polymerase II CTD (Pol II CTD) (proxy readout for transcription), H3K4m3, H3K27Ac, and histone H3 levels at indicated organ-specific genes using Matrix ChIP-qPCR. ChIP DNA was analyzed by qPCR, expressed as a fraction of input. In-house PCRCrunch software tool was used to download, analyze, and plot results. Data represent mean±SEM (n=4 for each frozen organ) expressed as a fraction of input. *Syn1*, brain-specific gene; *Tnnt2*, heart-specific gene; *Fxyd2*, kidney-specific gene; *Alb*, liver-specific gene; *Sftpa1*, lung-specific gene; and *Tnnc2*, muscle-specific gene.

### A high-throughput system for extracting soluble chromatin, DNA, and RNA from FFPE tissue blocks (Fig.2)

We used mouse brain, heart, kidney, liver, lung, and muscle FFPE blocks to develop and optimize the procedure for extracting analytes from FFPEs **(Fig.2A).** Standard formalin fixation (10% formalin for 24hrs) and paraffin embedding procedure was used to generate mouse organ FFPE blocks **(Fig.2B).** A microtome was used to cut FFPE slices. We began optimizing the methods with a standard protocol to deparaffinize FFPE curls with xylene, ethanol-H_2_O rehydration, and heat retrieval (23) followed by ultrasound shearing of soluble chromatin in PIXUL. The quality of soluble chromatin was assessed in Matrix ChIP using antibody to RNA polymerase II (Pol II) with IgG used as a background control, with the expectation that Pol II levels will be highest at genes expressed in a given organ. Primers were designed to genes specifically expressed in brain (*Syn1*), heart (*Tnnt2*), kidney (*Fxyd2*), liver (*Alb*), lung (*Sftpa1*), and muscle (*Tnnc2*) **(Fig.2C).** We found that input DNA generated from FFPE blocks contains PCR inhibitor(s). It has been reported that extracts from FFPEs can inhibit PCR due to small DNA degradation artifacts (46). To diminish this inhibition, we found that dilution of input samples by 20-fold minimized the qPCR interference. We tested a range of specimen slice thicknesses from 5 to 20μm and found that, for all the organs, 5μm worked well in PIXUL-ChIP assays while preserving more FFPE material compared to 10μm **(Fig.S1A-B).** We found that for some organs the Pol II ChIP signal could be noisy but subtracting IgG ChIP background signal greatly improved the consistency of PIXUL-FFPE results, a correction that has been included in the protocol.

**Fig.2.**
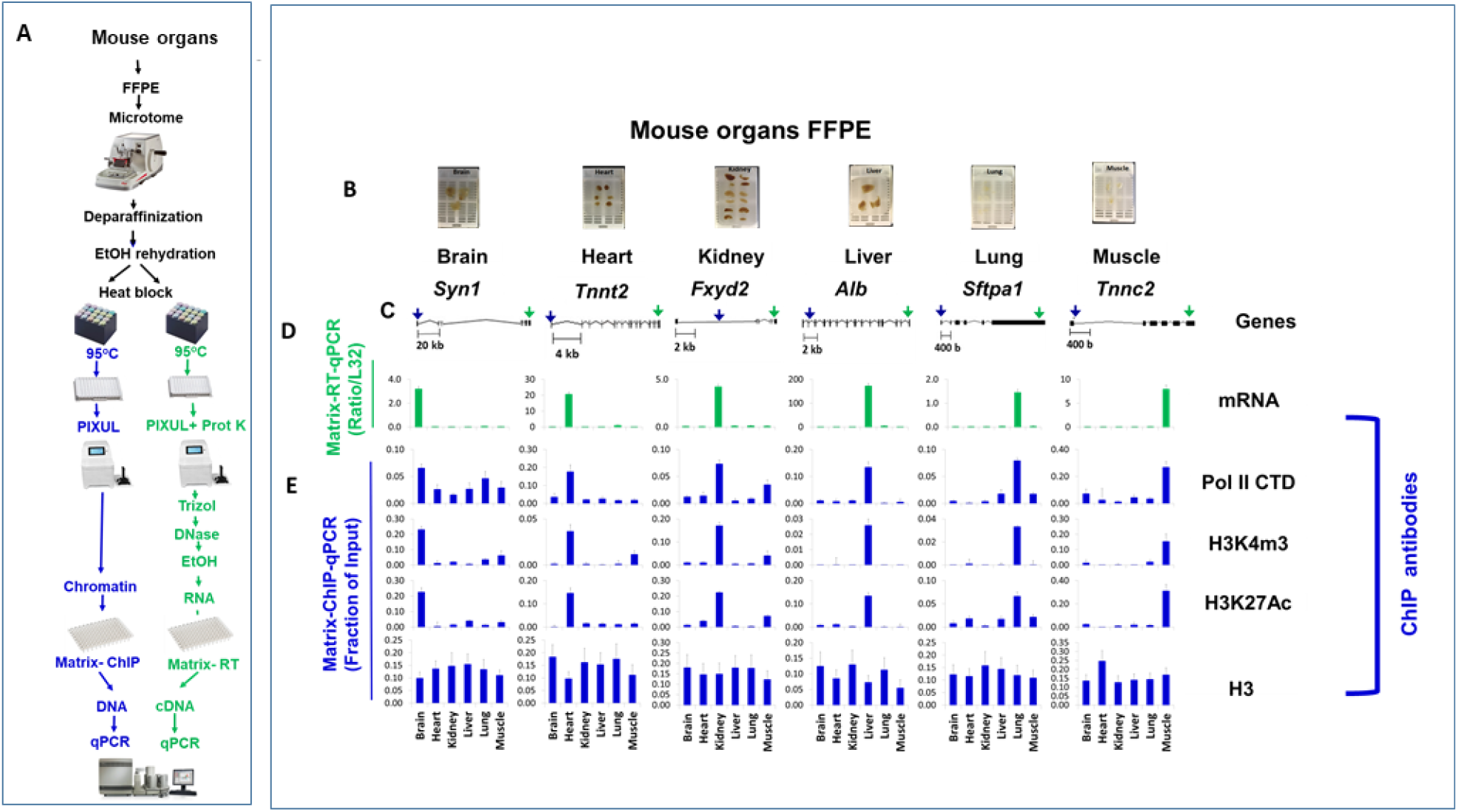
PIXUL-FFPE-Matrix-ChIP and mRNA analysis at mouse organ-specific genes. ***A.*** PIXUL-FFPE-Matrix-ChIP-qPCR and PIXUL-FFPE-mRNA-qPCR protocol. FFPE slices (5μm) were generated from FFPE blocks using a microtome, total 8 series for each organ. After deparaffinization, EtOH rehydration, and heat retrieval, samples were treated in PIXUL, and chromatin (*blue*) and RNA (*green*) were isolated*. **B.*** FFPE blocks of mouse brain, heart, kidney, liver, lung, and muscle. ***C.*** Cartoons of the genes. Blue arrows show positions of qPCR primers used in ChIP-qPCR and green arrows show positions of qPCR primers used in RT-qPCR. ***D.*** Extracted RNA was assayed in Matrix RT-qPCR using primers to 3’ ends of the genes. ***E.*** Extracted chromatin was analyzed in Matrix ChIP-qPCR using antibodies to Pol II CTD, H3K4m3, H3K27Ac, and histone H3. Mouse IgG was used for background subtraction. Inputs were diluted 20X to overcome PCR interference. Results (expressed as a fraction of input) represent mean±SEM (n=8 different FFPE extractions).

Xylene solvent has been traditionally used to deparaffinize FFPE blocks, including in epigenetics studies (23,47,48). However, it is toxic and, as such, hard to work with and less suitable for future PIXUL-FFPE automation. SafeClear is an organic, nonhazardous, nonflammable clearing and deparaffinization Xylene substitute (ThermoFisher). **Fig.S2A-B** show results of PIXUL-ChIP analysis using SafeClear compared to Xylene. Both solvents yield comparable tissue-specific patterns of permissive histone marks, H3K27Ac and H3K4m3, at 5’ ends of organ-specific genes, similar to Pol II profiles **(Fig.S2B).** As expected, histone H3 profiles were the same for each organ. These results demonstrate that both solvents perform similarly; therefore, SafeClear is used to deparaffinize blocks in the PIXUL-FFPE workflow.

Heating, typically at 95°C, is considered a critical step for retrieval of analytes from FFPE samples (49,50). And yet, heating can accelerate analyte degradation. Therefore, we tested temperatures lower than the standard 95°C for soluble chromatin retrieval. We compared chromatin extraction from rehydrated FFPEs at 95°C, 85°C, and 75°C. Samples were transferred to 96-well plates, treated with ultrasound in PIXUL, and analyzed in Matrix ChIP **(Fig.S3A).** These experiments showed that incubation at 95°C was more efficient than 85°C **(Fig.S3B)** and 75°C (not shown). We chose to use 95°C for heat retrieval of chromatin. Duration of incubation at 95°C was also important. We found that for ChIP analysis of extracted chromatin (Pol II, H3K4m3, H3K27Ac, and H3) 20min was better than 10min **(Fig.S4A-B)** while 5min yielded poor results.

We wondered if deparaffinization using SafeClear at higher temperature would improve the efficiency of chromatin extraction. We compared standard deparaffinization at room temperature (10min) which includes manual homogenization to 50°C (3min) without homogenization and found similar results in Matrix ChIP (Pol II, H3K4m3, H3K27Ac and H3) **(Fig.S5A-B).** We chose to carry out deparaffinization at 50°C to save time and eliminate the need for initial homogenization via pipette tip.

Some protocols include RNase treatment to increase chromatin recovery from rehydrated FFPEs (23,25,51). To test this, we incubated samples with or without RNase A (10μg/mL) for 30min at room temperature before the 95°C heating step. We found that application of RNase to rehydrated FFPEs did not make a difference in Matrix ChIP results (Pol II, H3K4m3, H3K27Ac and H3) **(Fig.S6A-B)** in our protocol. We chose not to include RNase treatment.

PIXUL uses polystyrene (PS) plates (27,28). For protocols that use high temperature incubation, 96-well polypropylene (PP) plates would be more suitable (PP is more heat resistant and has lower unspecific binding background). **Fig.S7** demonstrates that using 96-well PP plates in the PIXUL-FFPE workflow yields ChIP-qPCR results similar to those using 96-well PS plates (compare to **Fig.S6**). **Fig.S8** shows that substituting test tubes with 96-well PP plates in the heating step and then using the same plate for sonication in PIXUL yield the same ChIP-qPCR outcomes. These results demonstrate that using 96-well PP plates increases throughput of PIXUL procedure and simplifies workflow.

FFPE blocks can be stored at room temperature for several years (23,52). We found that soluble chromatin retrieved from 1.5-year-old FFPE blocks of mouse organs yielded Matrix ChIP results indistinguishable to those results obtained earlier from the same freshly prepared FFPE blocks **(Fig.S9A-B).** The blocks used in all the NGS studies were 1.5 years old (below).

RNA isolated from FFPEs can be effectively used in transcriptomic studies (53,54). We modified the PIXUL chromatin protocol for extraction of RNA from FFPEs. After incubation in extraction buffer at 95°C for 20 min, samples were transferred to 96-well plates and treated with ultrasound in PIXUL. Samples were transferred to 1.5mL tubes for standard TRizol RNA isolation protocol (55) followed by DNase treatment to eliminate residual genomic DNA. Finally, RNA was cleaned up either by EtOH precipitation or by Zymo clean up column **(Table S2).**

**Fig.2A** outlines final protocols of chromatin and RNA extraction from mouse brain, heart, kidney, liver, lung, and muscle FFPEs (**Fig.2B**) that were analyzed for transcript levels and epigenetic marks (**Fig.2C**). As in case of frozen organs (**Fig.1**), Matrix-RT-qPCR (**Fig.2D**) and Matrix-ChIP-qPCR (**Fig.2E**) data for organ-specific genes show that mRNA expression matches the pattern of Pol II and permissive histone marks (H3K4m3 and H3K27Ac) (**Fig.2D-E**). These comparisons serve to validate PIXUL-FFPE protocols.

The detailed workflows for extraction of RNA, DNA, and chromatin are shown in **Fig.S10** for fresh frozen tissues and in **Fig.S11** for FFPEs. These protocols were next adapted for next generation sequencing.

### PIXUL-Matrix-RNA-seq analysis of FFPE and frozen mouse tissues

RNA extracted with PIXUL from mouse FFPE (1.5-year-old) and frozen organs were used to generate sequencing libraries (**Fig.3A**). Correlation scatterplots (27) (deepTools) of FFPE vs frozen tissue BAM data showed Spearman correlation coefficients as follows: brain 0.91; heart 0.83; kidney 0.87; and liver 0.85 (**Fig.S13A**). IGV transcript snapshots shown in **Fig.3B** recapitulates results of RT-qPCR analysis of organ-specific transcripts in frozen (**Fig.1D**) and FFPE (**Fig.2D**) tissues. Housekeeping genes *Actb* and *Rpl32* are expressed in all four organs. The third intron of *Rpl32* gene encodes small nucleolar RNA, *Snord68,* that is also expressed in all four organs. Results of RNA-seq analysis further validate our PIXUL-FFPE RNA extraction method, as high-quality RNA can be extracted from FFPE specimens that were stored at room temperature for a long time.

**Fig.3.**
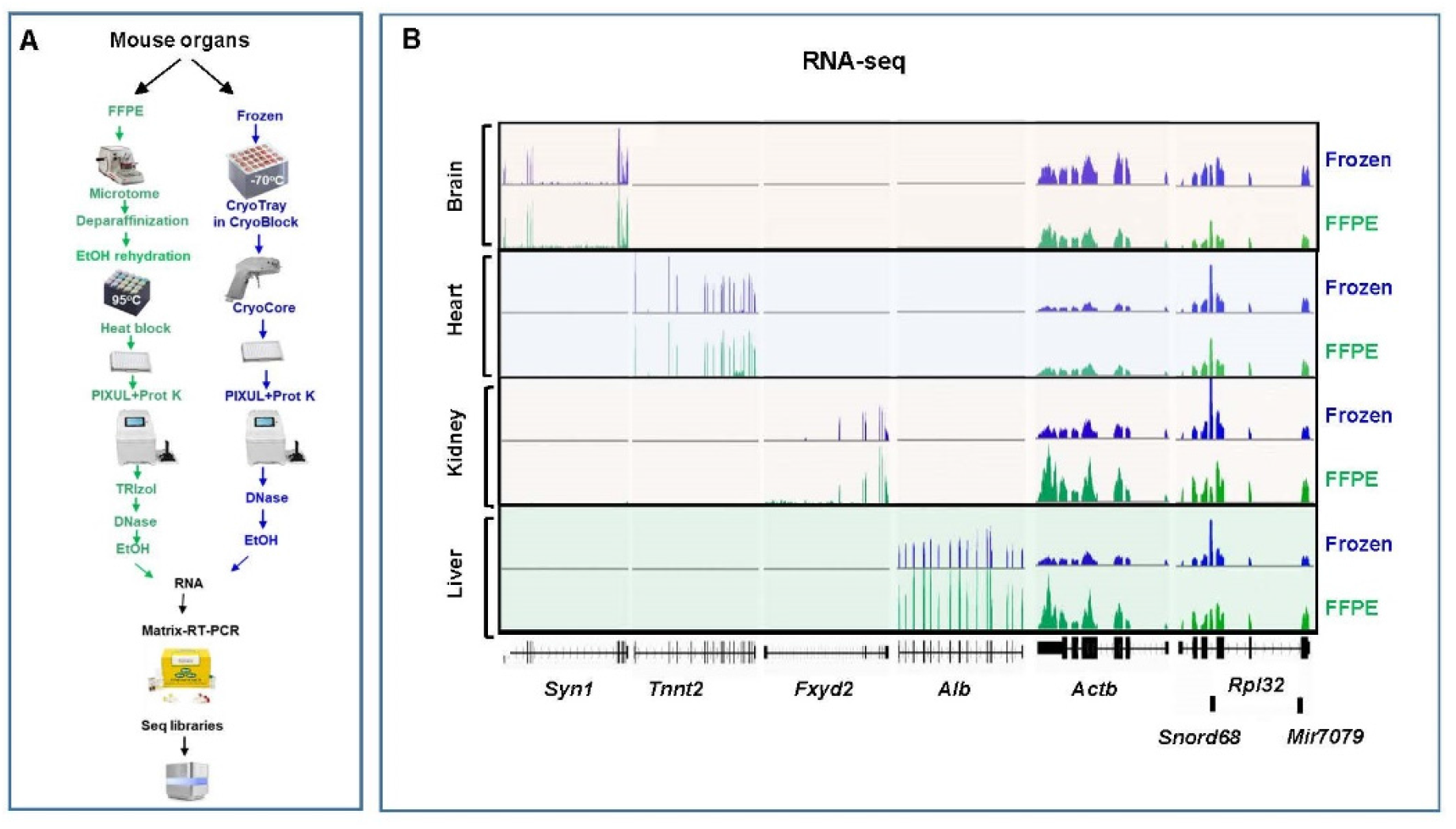
RNA-seq of FFPE vs. frozen mouse organs tissues analyzed by PIXUL-Matrix-RNA-seq. ***A.*** RNA was isolated from frozen (Fig.1) and FFPE (Fig.2) mouse tissues. Libraries were prepared and sequenced. Fastq files were downloaded from Illumina Basespace. Pre-processing was performed using fastp, and the quality of the runs was validated using fastqc (37). The pre-processed Fastq files were then aligned to the mouse mm10 genome and converted to BAM files using HISAT2 (paired-end library, reverse correspondence of read to transcript) (38). ***B.*** Transcript sequencing reads profile snapshots were generated from BAM files using Integrative Genomics Viewer (IGV) (132) of organ-specific RNAs; brain: *Syn1*; heart: *Tnnt2*; kidney: *Fxyd2*; liver: *Alb*; and housekeeping: *Actb* and *Rpl32*. Snord68 and Mir7079 are small nucleolar and micro RNAs respectively.

### Methylated RNA immunoprecipitation (PIXUL-Matrix-MeRIP)-seq analysis of FFPE and frozen mouse tissues

RNA modifications have been known for decades (56,57). With the introduction of MeRIP the number of studies to examine mRNA base modifications have increased, as they are thought to control transcripts’ translation and processing, a regulation called epitranscriptomics. m6A is one of the most common, reversible epitranscriptomic modifications (58,59). We have previously developed methylated DNA immunoprecipitation assay in 96-well plates (Matrix-MeDIP) (31) which we adapted here for MeRIP. RNA isolated from fresh frozen and FFPE tissues was further fragmented in PIXUL and immunoprecipitated with anti-m6A antibody bound to walls of 96-well plate via Protein A. After washes, MeRIP RNA was eluted and used to generate sequencing libraries (**Fig.4A**). Correlation scatterplots of FFPE vs frozen tissue BAM data showed Spearman correlation coefficient as follows: brain 0.85; heart 0.53; kidney 0.65; and liver 0.56 (**Fig.S13B**). **Fig.4B** shows the IGV m6A MeRIP-seq snapshot for the mouse brain, heart, kidney, and liver for organ-specific and housekeeping genes. MeRIP-seq looks very similar to RNA-seq except that there is no signal for the small nuclear RNA *Snord68*, suggesting that this RNA is not m6A modified. We found several other small nucleolar RNAs (snoRNAs) that are likewise not m6A modified, some of which are shown in **Fig.4C**. Note that the long non-coding RNA host *Snhg12* gene RNA is not expressed but the four different snoRNA, found in each one of its four introns, are not m6A modified (**Fig.4C**). To our knowledge, the lack of m6A base modification in small nucleolar RNAs has not been previously reported.

**Fig.4.**
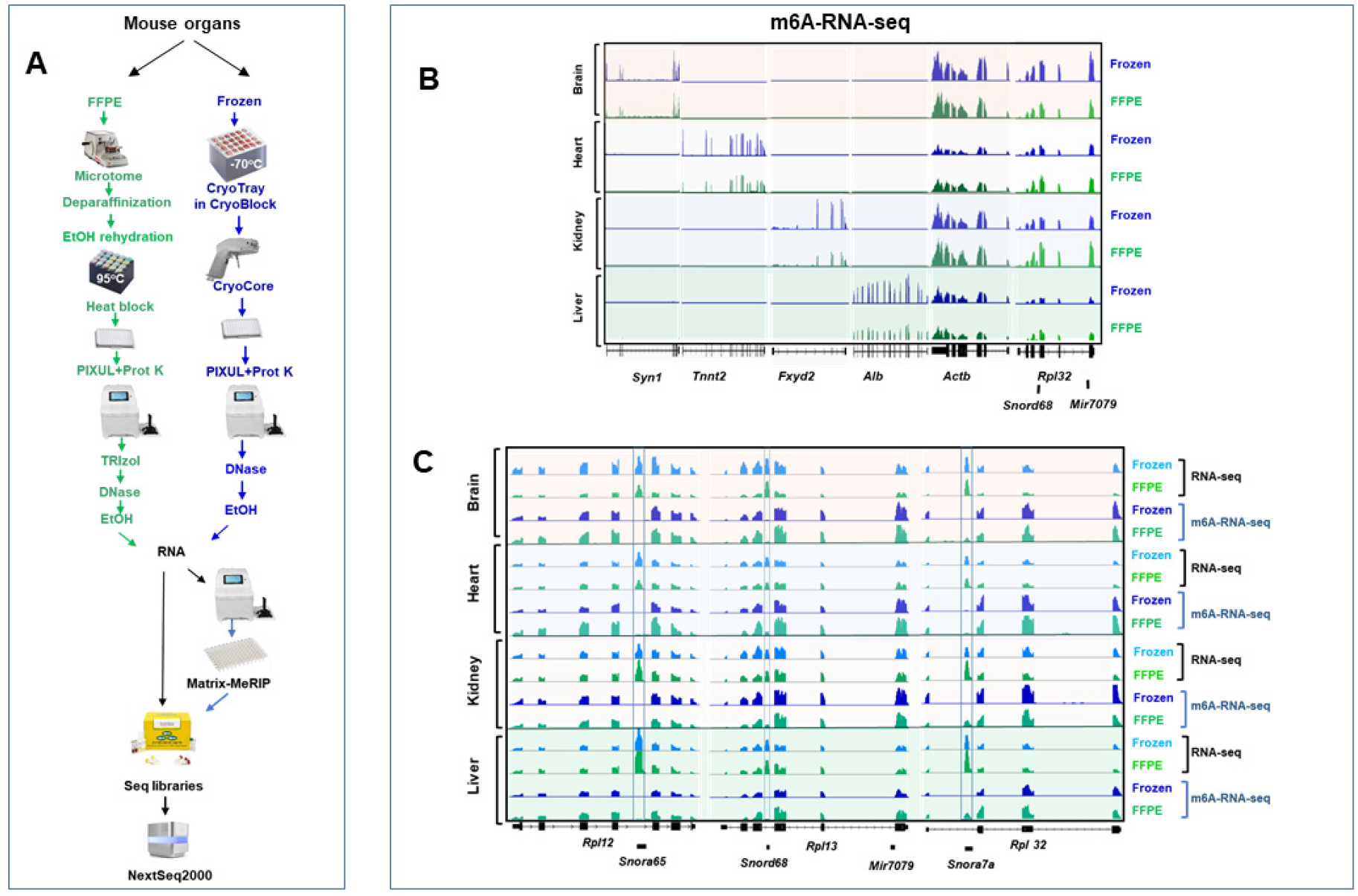
Methylated RNA (m6A)-seq FFPE vs. frozen mouse organ tissues analyzed by PIXUL-Matrix-methylated RNA immunoprecipitation (MeRIP). ***A.*** RNA was isolated from frozen (Fig.1) and FFPE (Fig.2) mouse tissues and was fragmented (PIXUL). Anti-m6A antibody in Matrix-MeRIP was used to immunoprecipitate methylated RNA. Libraries were prepared from methylated RNAs and sequenced. Fastq files were downloaded from Illumina Basespace. Pre-processing was performed using fastp, and the quality of the runs was validated using fastqc (37). The pre-processed Fastq files were then aligned to the mouse mm10 genome and converted to BAM files using HISAT2 (paired-end library, reverse correspondence of read to transcript) (38). ***B.*** Methylated RNA snapshots were generated from BAM files using Integrative Genomics Viewer (IGV) (132) of organ-specific RNAs; brain: *Syn1*; heart: *Tnnt2*; kidney: *Fxyd2*; liver: *Alb*; and housekeeping: *Actb* and *Rpl32*. Snord68 and Mir7079 are small nucleolar and micro RNAs respectively. Note that Snord68 is not m6A methylated. ***C.*** IGV snapshot demonstrating that snoRNAs are not m6A methylated. Note that the host gene lincRNA *Snhg12* RNA (133) is not expressed but the four non-m6A-methylated snoRNAs each found in one of the four introns are.

Small nucleolar RNAs (snoRNAs) belong to a well-characterized family of non-coding RNAs that play a role in rRNA maturation and also target tRNAs and mRNAs (60-62). Human snoRNA genes are mostly found within introns of mRNA or long noncoding RNA genes and are spliced out from the precursor RNA. N6-methylation of adenosine (m6A) in mRNA is co-transcriptional (63), but it has been shown that adenosine in introns have a small chance of being methylated compared to adenosine in exons (59). While mature mRNAs are exported to the cytoplasm for translation, snoRNAs are retained in the nucleus (64). Lack of m6A may prevent snoRNA cytoplasmic export.

### FFPE and frozen mouse tissues PIXUL-Matrix-ChIP-seq (Figs.5-7)

We used PIXUL-isolated chromatin from mouse FFPEs (1.5years old) and frozen organs in Matrix ChIP using antibodies to H3K27Ac **(Fig.5**), H3K4m3 (**Fig.6**), and RNA polymerase II (Pol II) (**Fig.7**). Scatterplots of FFPE vs frozen tissue BAM data showed good agreement with Spearman correlation coefficient as follows: H3K27Ac – brain 0.75; heart 0.82; kidney 0.80; and liver 0.79; (**Fig. S13C**); H3K4m3 – brain 0.76; heart 0.76; kidney 0.73; and liver 0.64 (**Fig.S13D**); Pol II CTD– brain 0.77; heart 0.74; kidney 0.76; and liver 0.69 (**Fig.S13E**). IGV snapshots shown in **Fig.5B** (H3K27Ac), **Fig.6B** (H3K4m3 and **Fig.7B** (Pol II) recapitulate frozen (**Fig.1D**) and FFPE (**Fig.2D**) ChIP-qPCR results at loci that encode organ-specific transcripts. Results of ChIP-seq analysis serve to validate the PIXUL-FFPE chromatin extraction method and demonstrate the usefulness of FFPE specimens in epigenetic analysis even after long-term storage at room temperature.

**Fig.5.**
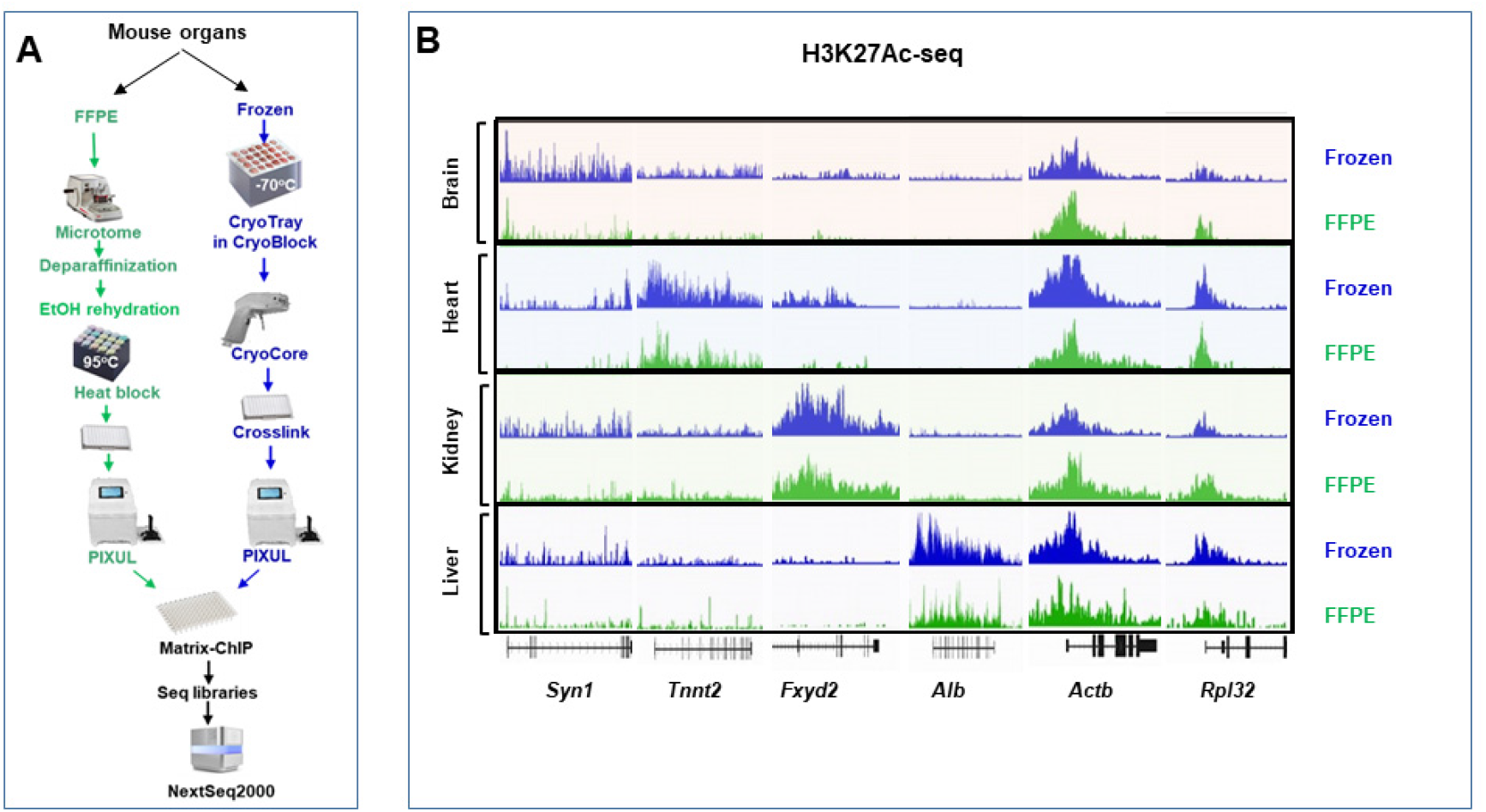
H3K27Ac-seq FFPE vs. Frozen analyzed by PIXUL-Matrix-ChIP-seq at mouse organ-specific and housekeeping genes. ***A.*** Chromatin was isolated from frozen (Fig.1) and FFPE (Fig.2) mouse tissues, and anti-H3K27Ac antibody was used in Matrix-ChIP. Libraries were prepared and sequenced. Fastq files were downloaded from Illumina Basespace. Pre-processing was performed using fastp, and the quality of the runs was validated using fastqc (37). The pre-processed Fastq files were then aligned to the mouse mm10 genome and converted to BAM files using Bowtie2 (134). Next, bamCompare (135) was used to normalize the antibody (treatment) files to their corresponding input (control) files (bin size = 10, scaling method = read count, comparison method = scaled value of treatment file), resulting in bigwig files. ***B.*** Snapshot was generated from bigwig files using Integrative Genomics Viewer (IGV) (132) of organ-specific H3K27Ac mark; brain: *Syn1*; heart: *Tnnt2*; kidney: *Fxyd2*; liver, *Alb*; and housekeeping: *Actb* and *Rpl32* genes.

**Fig.6.**
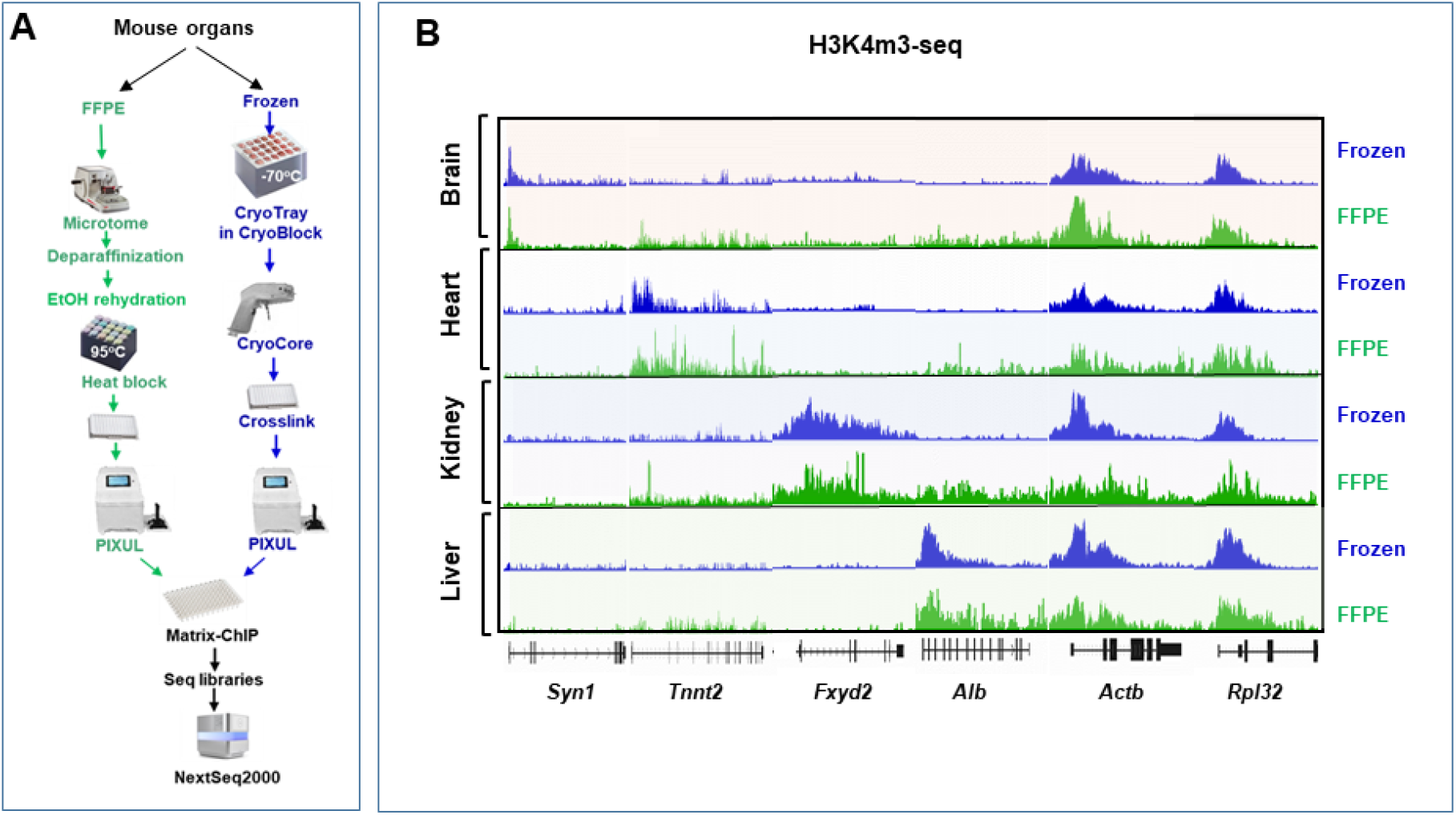
H3K4m3-seq FFPE vs. Frozen analyzed by PIXUL-Matrix-ChIP-seq at mouse organ-specific and housekeeping genes. ***A.*** Chromatin was isolated from frozen (Fig.1) and FFPE (Fig.2) mouse tissues, and anti-H3K4m3 antibody was used in Matrix-ChIP. Libraries were prepared and sequenced. Fastq files were downloaded from Illumina Basespace. Pre-processing was performed using fastp, and the quality of the runs was validated using fastqc (37). The pre-processed Fastq files were then aligned to the mouse mm10 genome and converted to BAM files using Bowtie2 (134). Next, bamCompare (135) was used to normalize the antibody (treatment) files to their corresponding input (control) files (bin size = 10, scaling method = read count, comparison method = scaled value of treatment file), resulting in bigwig files. ***B.*** Snapshot was generated from bigwig files using Integrative Genomics Viewer (IGV) (132) of organ-specific H3K4m3 mark; brain: *Syn1*; heart: *Tnnt2*; kidney: *Fxyd2*; liver, *Alb*; and housekeeping: *Actb* and *Rpl32* genes.

**Fig.7.**
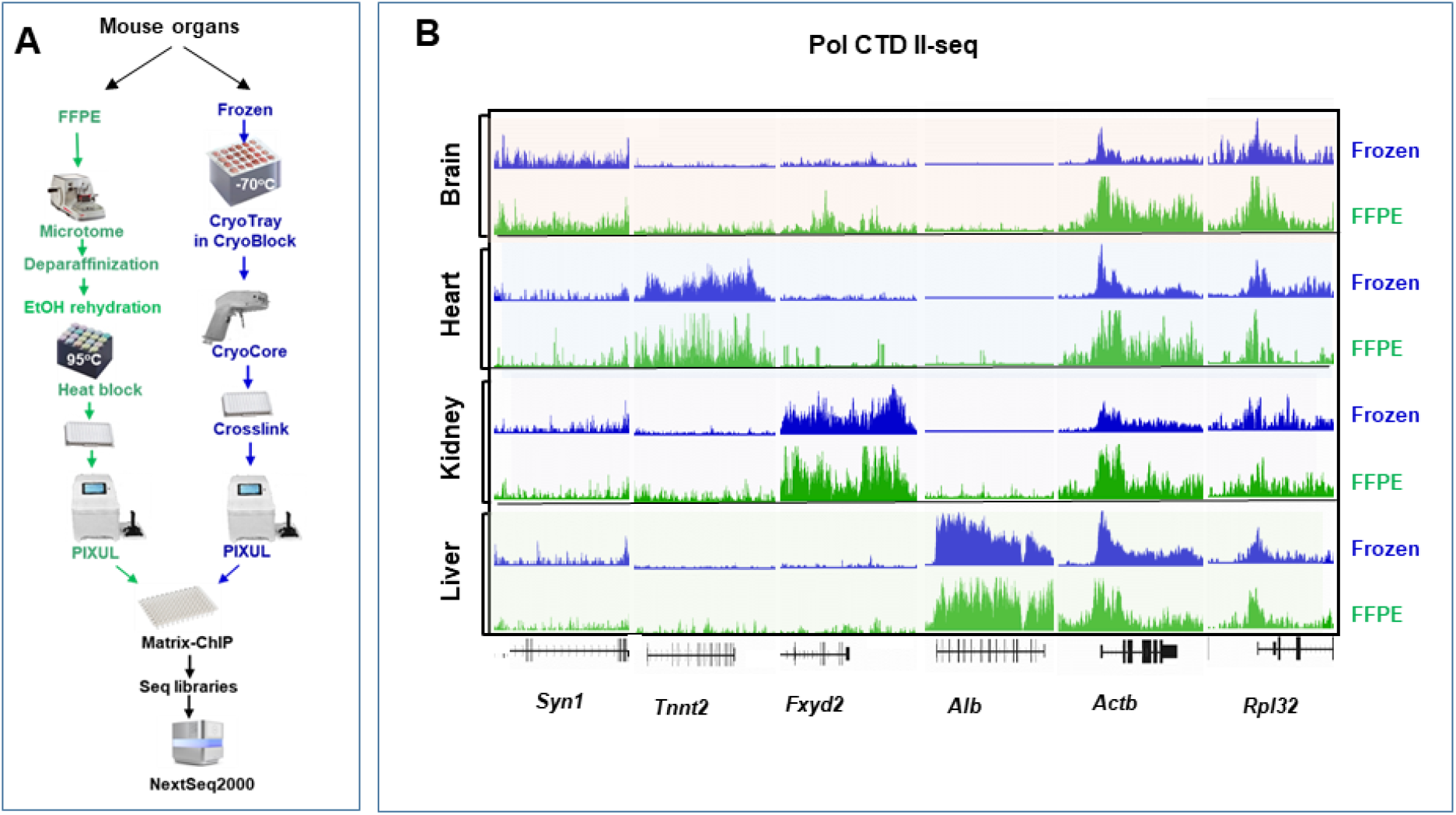
Pol II-seq FFPE vs. Frozen analyzed by PIXUL-Matrix-ChIP-seq at mouse organ-specific and housekeeping genes. ***A.*** Chromatin was isolated from frozen (Fig.1) and FFPE (Fig.2) mouse tissues, and anti-Pol II CTD antibody was used in Matrix-ChIP. Libraries were prepared and sequenced. Fastq files were downloaded from Illumina Basespace. Pre-processing was performed using fastp, and the quality of the runs was validated using fastqc (37). The pre-processed Fastq files were then aligned to the mouse mm10 genome and converted to BAM files using Bowtie2 (134). Next, bamCompare (135) was used to normalize the antibody (treatment) files to their corresponding input (control) files (bin size = 10, scaling method = read count, comparison method = scaled value of treatment file), resulting in bigwig files. ***B.*** Snapshot was generated from bigwig files using Integrative Genomics Viewer (IGV) (132) of organ-specific Pol II mark; brain: *Syn1*; heart: *Tnnt2*; kidney: *Fxyd2*; liver, *Alb*; and housekeeping: *Actb* and *Rpl32* genes.

### PIXUL-Matrix-MeDIP-seq analysis of FFPE and frozen mouse tissues

DNA methylation is the best studied epigenetic modification, and DNA methylation clinical biomarkers are used to stratify patients for cancer treatment (65-67). There are several DNA modifications. Methylation of the 5th position of cytosine (5mC) is the most common DNA modification (68). We adapted PIXUL protocols to extract DNA from FFPEs as follows (**Fig.8A**). Scatterplots of FFPE vs frozen tissue BAM data showed excellent agreement with Spearman correlation coefficient as follows: brain 0.89; heart 0.92; kidney 0.87; and liver 0.90 (**Fig.S13F**). Sequencing analysis reveals alternating DNA methylation pattern at 4.5S RNA loci corresponding to CpG islands (**Fig.8B**). Analysis of our RNA-seq, H3K4m3-seq, H3K27Ac-seq, and Pol II-seq data corresponding to 4.5S loci uncovered peaks of these marks in regions that are not methylated (**Fig.S14)**, suggesting that these permissive marks and 5mC are mutually exclusive (69). These data also show that like snoRNAs (**Fig.4**) 4.5S RNA appears not to be m6A modified. These Matrix-MeDIP-seq data validate our PIXUL-based DNA isolation method from FFPEs and demonstrate high quality of isolated DNA.

**Fig.8.**
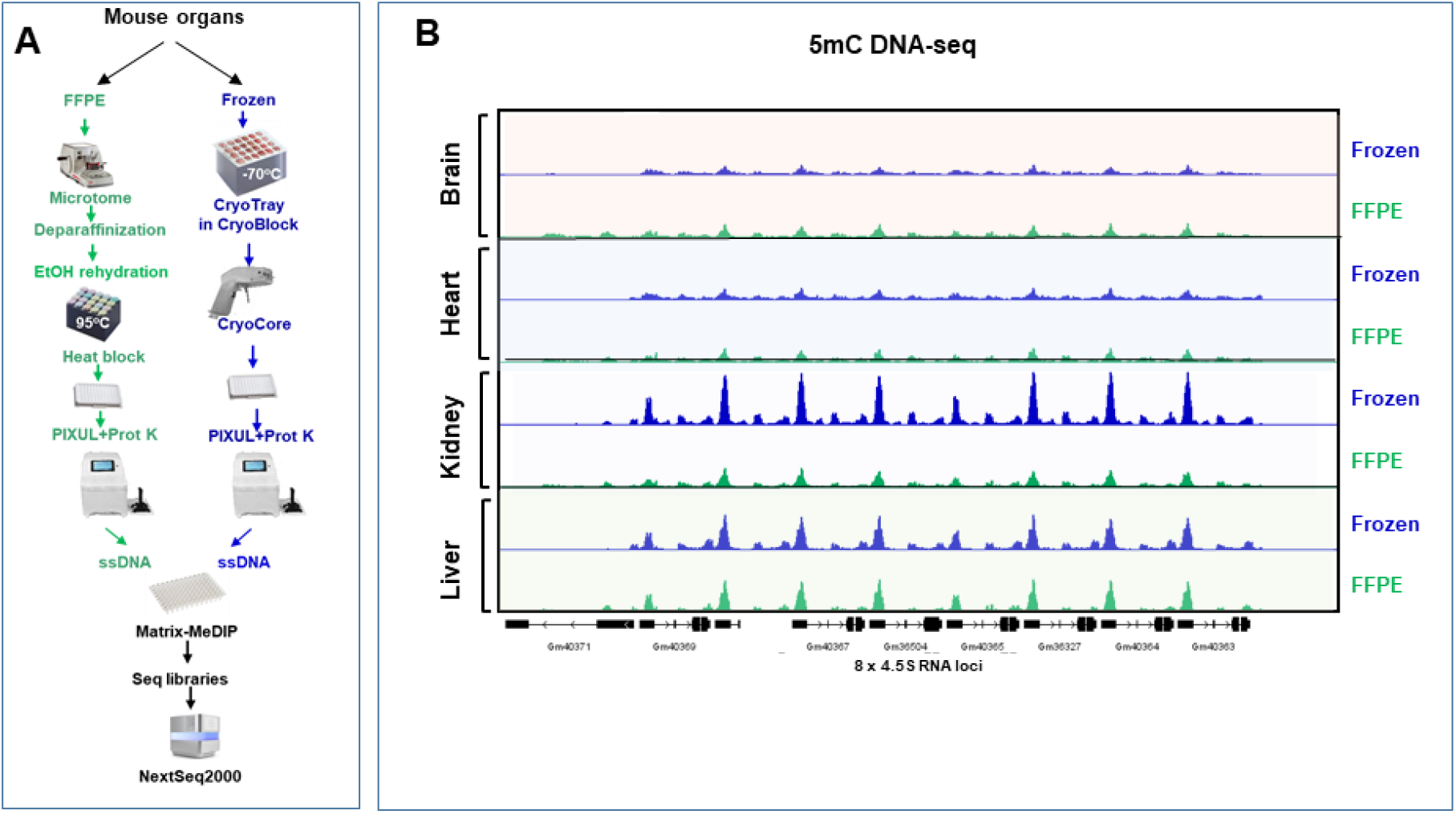
MeDIP-seq of FFPE vs. Frozen tissue analyzed by PIXUL-Matrix-MeDIP at 4.5S RNA mouse loci. ***A.*** PIXUL-based protocol was used to isolate DNA from FFPE and frozen mouse organs. Matrix-MeDIP was done using 5mC antibody (31). Libraries were prepared using ssDNA kit (xGen ssDNA & Low-Input DNA Library Prep Kit from IDT) and sequenced. Fastq files were downloaded from Illumina Basespace. Pre-processing was performed using fastp, and the quality of the runs was validated using fastqc (37). The pre-processed Fastq files were then aligned to the mouse mm10 genome and converted to BAM files using Bowtie2 (134). Next, bamCompare (135) was used to normalize the antibody (treatment) files to their corresponding input (control) files (bin size = 10, scaling method = read count, comparison method = scaled value of treatment file), resulting in bigwig files. ***B.*** Snapshot was generated from bigwig files using Integrative Genomics Viewer (IGV) (132) at a tandem 4.5S RNA loci.

### PIXUL-LC-MS/MS in FFPE and frozen mouse tissues

Recently, significant progress has been made in extracting proteins from tissues, including FFPE, for proteomic analysis (20,70,71). Still, to the best of our knowledge, there are no studies that combine proteomic measurements with epigenomic, transcriptomic, and epitranscriptomic assays in the same biospecimens, let alone FFPEs. To mitigate sample bias and batch effects (72), our goal was to develop a 96-well format proteomic workflow aligned with all the other omics. (**Figs.9A and S13)**. Comparable numbers of proteins were identified and quantified in FFPE vs. frozen organs: brain 5267 vs. 5182; heart 5321 vs. 4978; kidney 5292 vs. 5162 and liver 5299 vs. 5076 (**Fig.9B**). There was a high correlation of protein abundances between FFPE and matched frozen organs: brain 0.969; heart 0.947; kidney 0.957; and liver 0.953 (Pearson R^2^) (**Fig.9C**). In agreement with large-scale proteomic studies (73) Venn diagrams illustrate that the number of organ-specific proteins ranged between 24-90 or 0.5-1.9% of the total number of a given organ **(Fig.9D).** These results demonstrate that proteomic analysis can be combined with epigenomic, trancriptomics and epitransciptomic assays from the same samples using the PIXUL.

**Fig.9.**
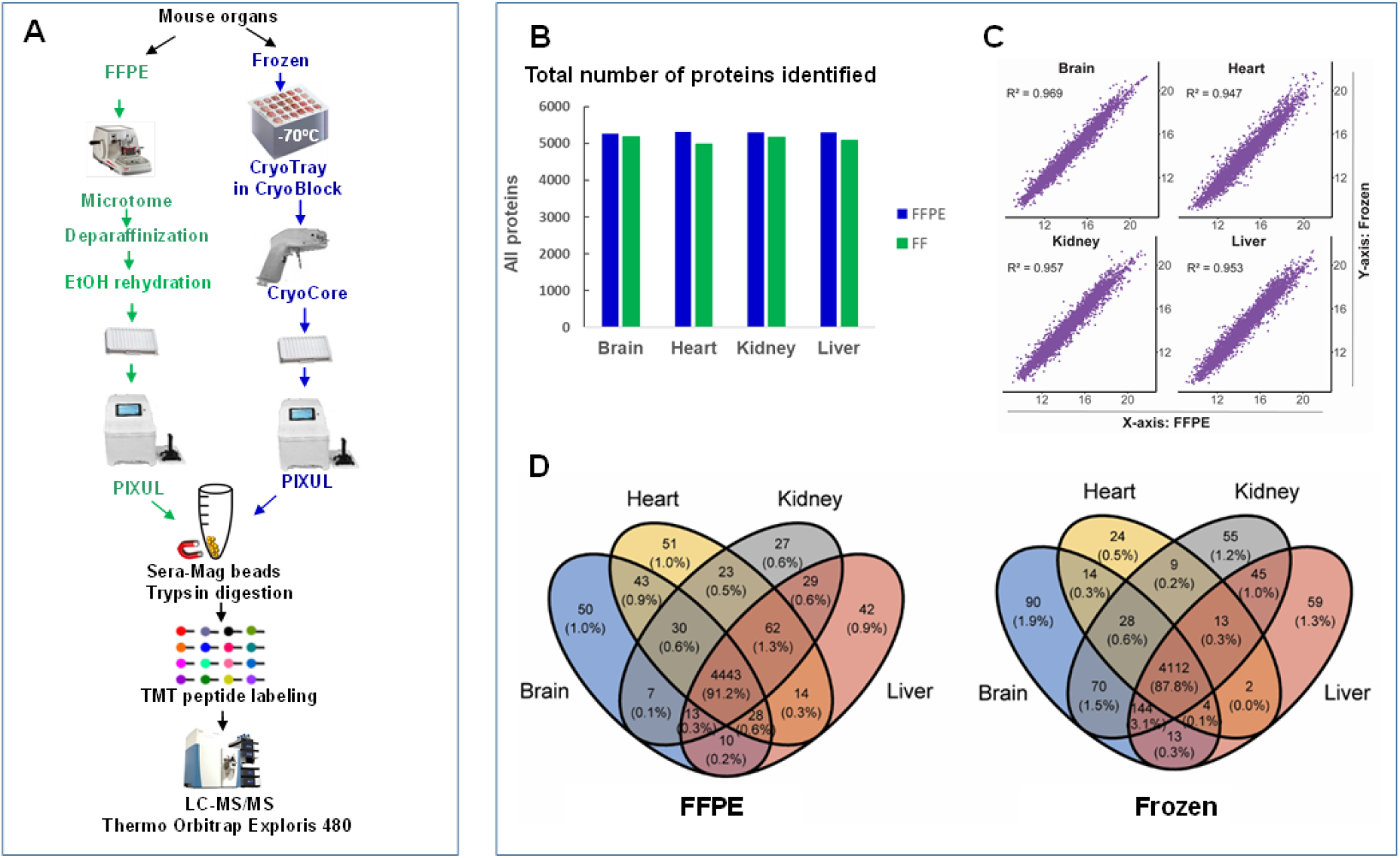
Protein extracted from FFPE and Frozen tissue analyzed by PIXUL-LC-MS/MS. ***A.*** PIXUL-based protocol was used to isolate proteins from matched FFPE and frozen mouse organs. CryoCore was used to sample frozen tissues and microtome to sample FFPEs. Proteins were extracted (Methods) using PIXUL and trypsin-digested using SP3 protocol, and TMT-labeled peptides were fractionated and analyzed by LC-MS/MS. ***B.*** Total number of proteins identified in organ from frozen and FFPE blocks. ***C.*** Scatter plots of comparative analysis of PIXUL-LC-MS/MS generated protein datasets from mouse organs frozen and FFPE blocks. ***D.*** Venn diagram comparing proteins identified across the four organs.

### The Segway algorithm produces annotations of the genome based on ChIP-seq and MeDIP-seq datasets

Segway (41) is a dynamic Bayesian network model that takes as input heterogeneous genome-wide measurements and coalesces them into a segmentation of the genome and an associated set of labels. Here we ran Segway across multi-tissue multi-omics profiles (ChIP-seq: H3K27Ac/H3K4m3/Pol II and MeDIP-seq: 5mC datasets) on FFPE and frozen separately, and we generated genome segmentations and annotations for mouse brain, heart, kidney, and liver tissues (**Fig.10A**). Genomes in each tissue were annotated with 15 labels, each representing a specific type of genomic function (e.g., promoters and enhancers). Specifically, we identified one segmentation label for FFPE and frozen samples each that is enriched for previously annotated super-enhancers (dbSUPER, (42)) (**Fig.10B**). Interestingly, Segway’s organ-specific annotation of super-enhancers agrees with previous analyses (42), suggesting that multi-omics data generated by PIXUL-Matrix-ChIP/MeDIP-seq can reveal organ-specific epigenomic regulation. This result also demonstrates that FFPE measurements alone can be used to capture organ-specific regulatory elements. We further derived a set of super-enhancers that are predicted to only occur in one organ and are identified in both FFPE and frozen samples (brain 3342; heart 1059; kidney 1642; and liver 3029, **Fig.10C**). Thus, in addition to re-identifying enhancers that are already annotated in dbSUPER, Segway is able to generate new hypotheses by predicting a set of new organ-specific super-enhancers based on our multi-omics profiles.

**Fig.10.**
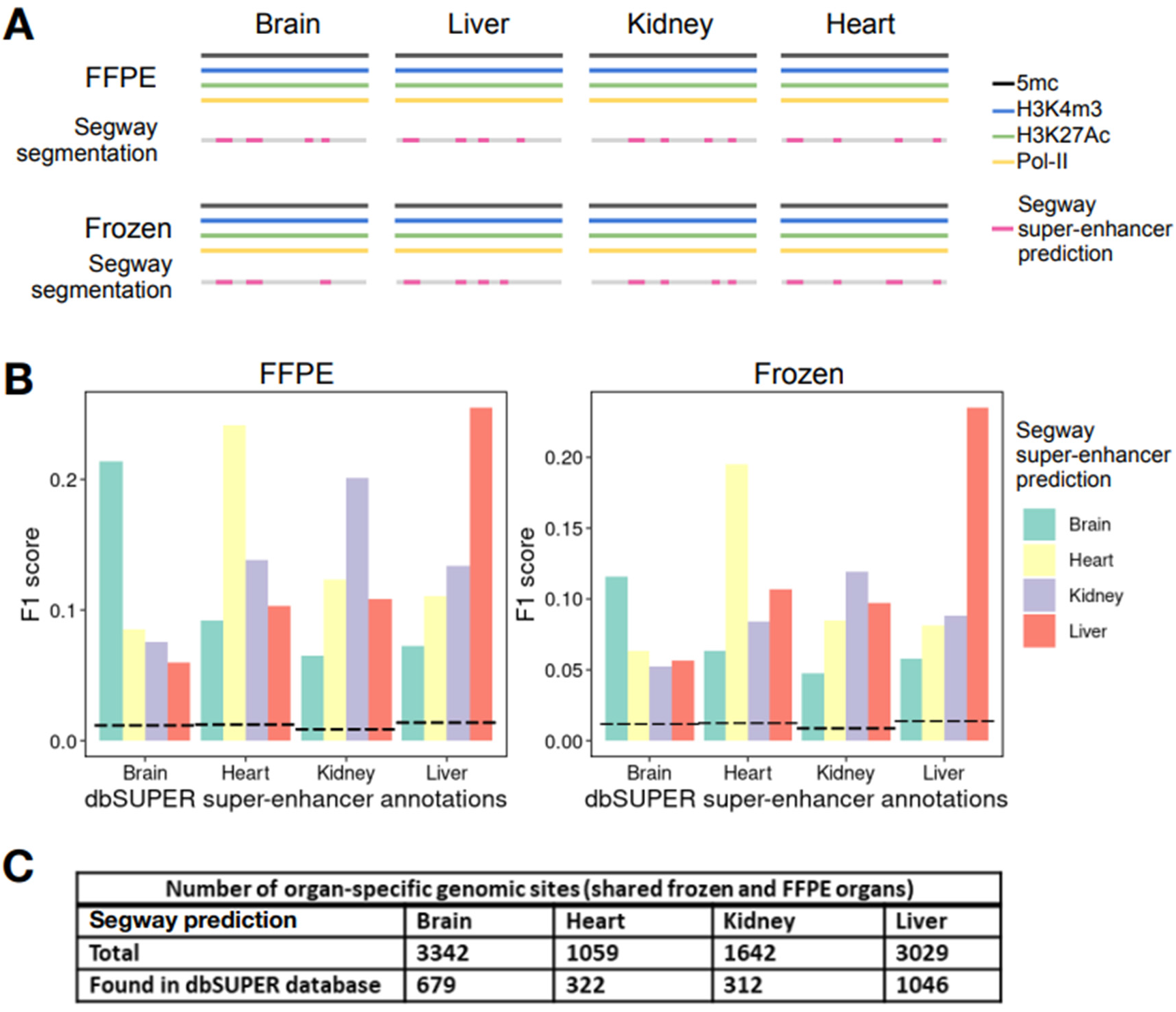
Tissue-specific super-enhancer prediction by Segway and comparison to dbSUPER’s tissue-specific super-enhancer annotations. ***A.*** Segway protocol. ChIP-seq: H3K27Ac/H3K4m3/Pol II and MeDIP-seq: 5mC datasets were used. A Segway model is trained on frozen and FFPE samples separately and annotates the four genomes into 15 types of genome segments, one of which are enriched for super-enhancers (*pink region*). ***B.*** For each organ, an F1 score based on precision and recall with respect to dbSUPER’s super-enhancer annotations was calculated. Dotted lines show F1 scores from a random baseline. dbSUPER’s tissue-specific super-enhancers were best captured (with the highest F1 score) by Segway’s annotation in the corresponding tissue compared to unrelated tissues. ***C.*** Numbers of predicted organ-specific super-enhancer sites shared in both frozen and FFPE organs and numbers of these sites also found in dbSUPER database.

### Predicting protein abundance using multi-omics profiles

Tissue protein and mRNA levels for a given gene are not well correlated (73-75). For instance, in the ageing kidney changes in protein levels can occur without changes in cognate mRNAs (76). Because there is a large discrepancy between transcriptomic and proteomics expression measurements (**Fig.11A**), we further investigated whether multi-omics profiles can provide a more accurate predictor of protein expression. To do that, we trained linear regressors based on different combinations of multi-omics profiles to predict protein expression levels across organs and samples. We found that protein expression is best predicted when using both DNA- and RNA-based predictors (**Fig.11B-C, Fig.S15**). In addition, compared to RNA expression, m6A RNA methylation level is a better indicator of protein expression (**Fig.11D, Fig.S16**). Furthermore, combining epitranscriptomic measurements with transcriptomics markedly improves protein expression prediction (**Fig.11E**). These findings suggest that epigenomic and epitranscriptomic assays provide complementary information to transcriptomics measurements in relation to protein quantities, demonstrating the potential of integrating multi-omics measurements to model proteomics profiles.

**Fig.11.**
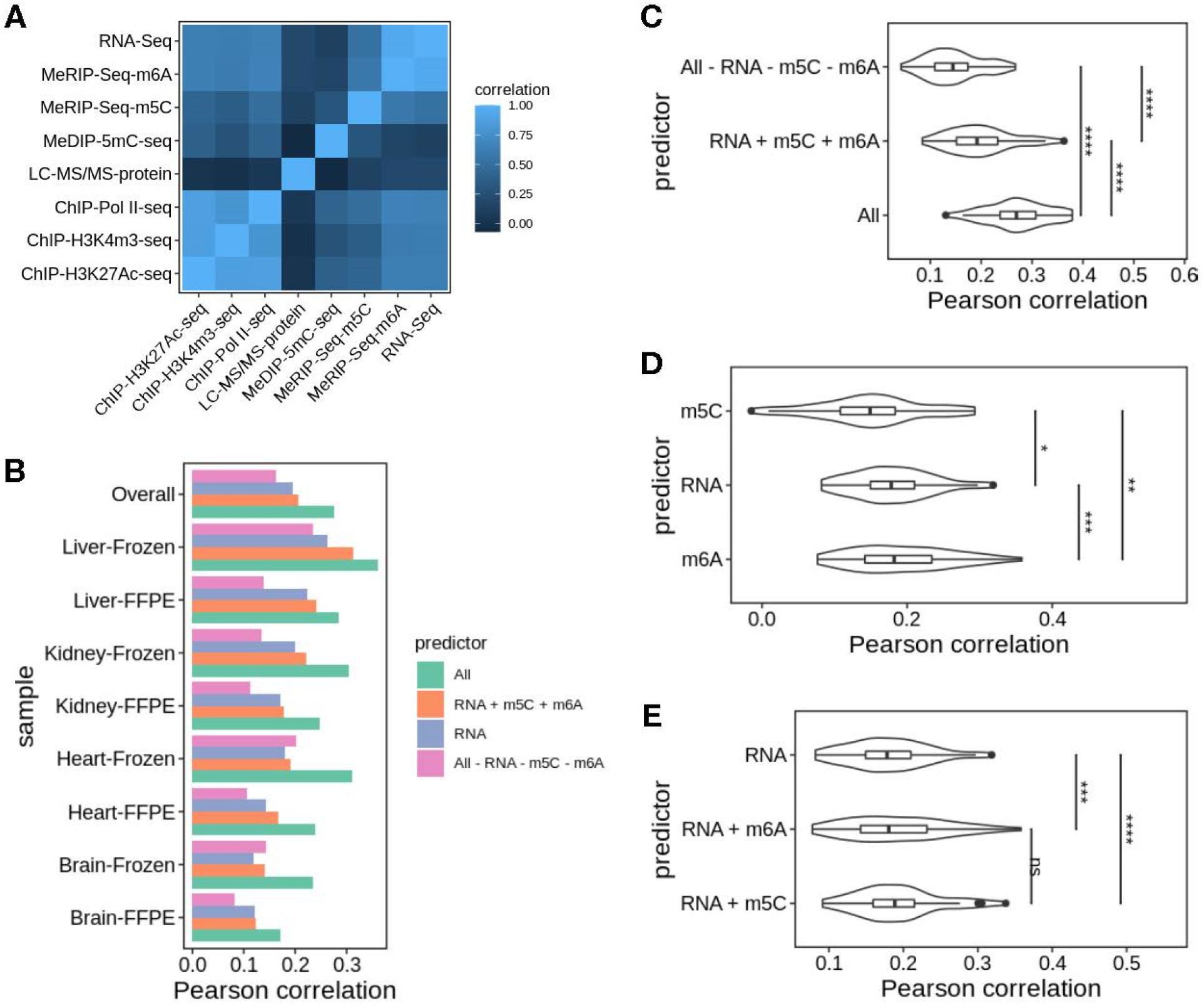
Protein expression prediction using multi-omics profiles. ***A.*** Pairwise Pearson correlation between observed multi-omics datasets profiles. ***B.*** Pearson correlation between predicted and observed protein expression, separated by organ and sample prep. Protein expression is predicted by several sets of multi-omics profiles: “All”: all other seven omics types in this study; “RNA + m5C + m6A”: transcriptomic and epitranscriptomic profiles; “RNA”: transcriptomics profiles; “All – RNA – m5C – m6A”: epigenomics profiles. ***C-E.*** Violin plots comparing the performance of protein expression prediction across different sets of predictors. Statistical significance is calculated through one-sided paired Wilcoxon signed-rank test on the Pearson correlation coefficients across cross-validation folds, organ, and sample prep. P-values were subjected to Benjamini-Hochberg correction.

## DISCUSSION

Advances in analytical methods provide new opportunities for profiling multiomes in cells and tissues. Still, progress in high throughput and efficient extraction of analytes from biospecimens has been slow, and multi-omics studies have been limited to selected cultures or tissues. To facilitate multiome analysis, we used frozen and FFPE blocks from mouse organs to develop PIXUL ultrasound-based 96-well-format methods to extract chromatin, DNA, RNA, and proteins from these biospecimens. We integrated these sample preparation protocols with downstream multi-omic analysis platforms, and in so doing, made these workflows, which we call MultiomicsTracks96, useful for high throughput multiome analysis of frozen and FFPE biospecimens.

### Application of ultrasound for analytes extraction and molecular assays

Ultrasound is increasingly being used in a variety of sample preparation and analytical applications. It is commonly used to fragment DNA, chromatin, and RNA for downstream analysis (27,77,78). Low energy ultrasound facilitates molecular interactions (79-81) and has been used in analytical applications including immunoassays (79) (for example, immunoprecipitation (82)) and for trypsin digestion (83). The use of sonication to extract substrates from insoluble material goes back nearly 60 years (84). The food science and technology sectors have been early adaptors of ultrasound to extract molecules (85-87), including oils (88). Ultrasound can also be used to facilitate emulsification, including paraffin wax (89). More recently, ultrasound has been used to extract DNA (90), RNA (53), chromatin (78), and proteins (83) from a variety of biospecimens, including FFPEs. Still, these methods have remained slow and low throughput. We have previously engineered a 96-well format PIXUL sonicator to treat samples in 96-well plates (27-29). Here we adapted PIXUL for extraction of chromatin, DNA, RNA, and protein from frozen and FFPE tissues for 96-well format multi-omics applications.

### FFPE tissue blocks for nucleic acids analysis

Formalin fixation and paraffin embedding is the most commonly used method to store and process biospecimens for histopathology. FFPE preserves tissue and cell morphology and architecture to allow histological interpretation (91). The use of formaldehyde for tissue fixation goes back to 1893 (91,92). In the 1940s, fixation chemistry research showed that formalin reacts with proteins through amino groups (93-95), and later it has been shown to cross-link to other moieties, including amino groups of DNA (95). For decades, FFPE blocks were used primarily for histopathology until the late 1980s, when DNA (96) and RNA (97) were first isolated from FFPEs and used in PCR (97-99). These early nucleic acid extraction methods introduced various approaches such as Proteinase K proteolytic treatment with heating (96,99), boiling (98), and sonication (100). Application of silica adsorption in the 1990s further improved the purity of nucleic acids extracted from FFPEs (101). In 1993, Shi et al. demonstrated that heat can be used to retrieve antigens from FFPE tissues, representing the first evidence that these archived tissues can be used for protein analysis (102-104). Although these early methods were slow, inefficient, and yielded variable results, this pioneering work represented a paradigm shift, demonstrating that FFPE blocks, traditionally used for histopathology, can be used in proteomic (21), genomic (48), transcriptomic (105), and epigenomic (106,107) applications.

Given the enormous potential of FFPEs in basic research and clinical applications (26), many companies have developed kits to extract DNA, RNA, or both from these biospecimens. These commercially available tools use Proteinase K, heating, sonication, and silica adsorption techniques developed by the early pioneers of the field (19). Comparative analysis revealed that no single kit was superior to the others, and all performed well in downstream next generation sequencing applications (NGS) (108-111) that showed high concordance with paired fresh frozen tissues (54,111,112). FFPE as old as 12 years can be efficiently used for NGS analysis (26). Our 96-well format PIXUL-Matrix method also demonstrated high RNA-seq concurrence between FFPE and frozen mouse organs tissues **(Fig.S13A).**

### FFPE blocks for proteomic analysis

In 2005, Hood et al. tested whether archived FFPE could be used for proteomic analysis and isolated peptides representing several hundred proteins from prostate cancer and benign prostate hyperplasia (113). Using paired fresh frozen and FFPE mouse liver tissues, they identified 776 and 684 unique proteins, respectively. This study demonstrated for the first time that FFPE can be used effectively to obtain reliable proteomic data (113). Subsequently, studies by Guo et al. (114) and Sprung et al. (115) demonstrated 83% to 92% overlap of proteins identified in pairs of fresh frozen and FFPE tissues. More recently, FFPEs were tested in large-scale proteomic analysis, taking advantage of either ultrasound (20,70) or pressure cycling technology (71). These latest studies demonstrated that FFPEs as old as 15-20 years can be used effectively for proteomic analysis even though there is some decrease in peptide identification compared to newer samples (70). Correlation of paired fresh frozen and FFPE tissues in these studies was high with *R*^2^> 0.9 (20,71). Our PIXUL-LC-MS/MS analysis yielded *R*^2^ ≥0.95 **(Fig.9C).**

### FFPE blocks for epigenetic analysis

Historically, compared to other analytes, extraction of soluble chromatin from FFPEs has been challenging, but recently there has been significant progress made using these biospecimens for epigenetic analysis (reviewed (23)). In 2010, Fanelli et al. reported an improved method for epigenomic analysis from FFPEs, which they termed PAT-ChIP (25). To isolate chromatin, they used sonication and controlled micrococcal nuclease (MNase) treatment. Later, to further increase efficiency of chromatin extraction, the same group introduced a heating step (1hr at +80°C) and eliminated MNase. The improved method was named enhanced PAT-ChIP (EPAT-ChIP) (51). In 2016, Cejas et al. described a method that uses limited Proteinase K treatment to extract chromatin from FFPEs for ChIP-seq analysis, FiT-ChIP (52). This method underperformed in H3K27Ac-seq analysis. To mitigate this shortcoming, the same group published another protocol called FiTAc-seq that eliminated Proteinase K and added a heating step (49). Other methods for epigenome analysis of FFPE from other labs followed: Chromo-EX PE (116), RCRA-ChIP-seq (117), and a Tn5 transposase-protein A-based FACT-seq (118). All these newer protocols include heating steps (23). For all the published methods, the time needed to prepare sequencing library ready-DNA ranged from 2-3 days for RCRA-ChIP-seq to 5 days for FACT-seq. There are also commercially available FFPE-ChIP-seq kits, including ChIP-IT FFPE from Active Motif and truChIP FFPE from Covaris. The latter is the only FFPE-ChIP protocol that uses sonication to emulsify paraffin. PAT/EPAT-ChIP and the ChIP-IT FFPE are the only methods that have been used in studies published by other researchers. Further, the number of FFPE ChIP citations is less than 20 (23) which may in part reflect the fact that these methods remain challenging. In contrast, the PIXUL-Matrix-ChIP FFPE method is simple and takes less time: starting from a tissue fragment it takes 1 day for qPCR-ready DNA and 1.5 days for sequencing library ready DNA **(Fig.S11, Table S5).**

### FFPE blocks for epitranscriptomic analysis

Until now, to the best of our knowledge, FFPE tissue blocks have not been used in epitranscriptomics studies. RNA methylation was first described by Littlefield and Dunn in 1958 (119-121). Decades of studies that followed demonstrated the ubiquitous nature of methylation in different RNA species in all kingdoms of life (122). Addition of methyl groups to nitrogen, carbon, and 2’OH moieties are by far the most common among the ∼170 known post-transcriptional RNA modifications (122-124). Messenger RNA methylation was first described in 1974 (56), but research in this area lagged behind studies of tRNA, rRNA, and snRNA modifications (124). Advances in genomic technologies stimulated studies to explore mRNA post-transcriptional modifications. In 2012, Dominissini et al. (123,124) and Meyer et al. (125) used, for the first time, MeRIP-seq to profile levels of m6A transcriptome-wide, research that paved the way for epitranscriptomics. m6A is the most common internal mRNA modification found in eukaryotic and single-cell organisms (123,124) and one of the best studied (58). The GLORI method allows single-base m6A mRNA transcriptome-wide analysis that uses deamination of unmethylated adenosines (126) and is the most recent addition to the epitranscriptomics tool box (58). The concurrence of GLORI-identified m6A sites with the more traditional methods, including MeRIP, is approximately 50% and, as such, the reliability of this costly technique will need to await further improvement (126). The regulatory role of m6A has been implicated in virtually every step of RNA-mediated gene expression, including splicing, nuclear export, and translation, to name just a few (58,63). The fundamental role of m6A in gene expression does not appear to be unique. Rather, its role seems to be shared with other RNA base modifications, even though their levels in mRNAs are 10-fold lower (e.g., m1A, m5C) (124). Thus, it is conceivable that the combination of mRNA base modifications constitutes an epitranscriptomic code. In this regard, our PIXUL-Matrix-MeRIP platform should greatly facilitate epitranscriptome profiling of a wide range of biospecimens, including FFPEs.

### High dimensional multi-omics

Single-ome datasets provide a biological signature limited to a lone intracellular tier (5). In such a mono-omic view, a network of relationships (edges) between molecules (nodes) can be constructed, but the ability to link single profiles to phenotypes is limited (3). In contrast, integration of multi-omic datasets aims to map the flow of information spanning multiple intracellular organization layers and, in so doing, define causality (1,2,9). As such, multi-omics is a systems biology approach to better understand biological processes and phenotypes in health and disease, advancing diagnosis and treatments (1-3,9). Still, multi-omics is a nascent discipline in which most studies have included only two omics datasets (1). For instance, in a 2022 review of the NIH multi-omics grant portfolio, which included metabolomics, only 7% and 2% of projects consisted of four and five omics approaches, respectively (127). Our study measures 8 different omic datasets, including, for the first time, epitranscriptome profiles. Further, our multidimensional survey covered several organs while comparing frozen and FFPE tissue blocks.

Despite great advances in analytical tools, integrating heterogeneous multi-omics datasets remains computationally challenging (1,2,9). Technical variation and sample batch effects render the integration of multi-dimensional datasets more difficult (9,72,128). To mitigate experimental biases, we developed protocols that use a single sample for all the bulk multiome assays. Additionally, we aimed to develop workflows to maximize the number of sample preparation steps shared by the different multi-omics assays.

### Bulk vs. single-cell multi-omics

The bulk multi-omic datasets report averages across all cells in an assayed solid or liquid biospecimen. As such, bulk multi-omics studies are an informative, practical, and cost-effective means to define differences between specimens, as is the case here where we compared multiomes of several mouse organs. Assays of tissue fragments or liquid biospecimens are well suited for large-scale studies (8,70,129). However, bulk multi-omics assays lack the necessary resolution to infer tissues’ cellular heterogeneity and may not detect signals from less abundant cell types. Missing this type of information could limit interpretation of integrated multi-omics data. In contrast, single-cell studies allow one to define tissue cellular heterogeneity and identify cell types. While initially focused on transcriptomics (single cell RNA-seq), more recent single cell studies have reported two (e.g., transcriptome and methylome) or even three (e.g., genome, transcriptome, and methylome) multi-omics datasets (130,131). These single-cell studies provide information about cell types, cell-cell communication, cell lineages, and regulatory networks. However, compared to bulk assays, single-cell measurements are plagued by greater technical noise, lower depths, complex experimental protocols, high cost, and impracticality for large-scale biospecimen studies required for longitudinal clinical studies (e.g., TopMed (8)). Here, we present same sample 8 omic datasets from four different organs, a degree of multiplicity not feasible for single-cell studies in the foreseeable future, let alone from FFPEs. When it comes to epigenetics, epitranscriptomics, or proteomics, only a few out of many known nucleic acids and protein modifications have been assayed thus far in multi-omics studies (1,127). In this regard, extending multiome measurements to include several molecular classes within any given omics data type (e.g., phospho- and glycoproteins for proteomics, or multiple DNA or RNA modifications) is quite feasible for bulk but not for single-cell assays. In sum, both MultiomicsTracks96 and single-cell methods have limitations that the other approach could mitigate. As such, bulk and single-cell assays provide complementary means to better understand the multiomes that define phenotypes. (130).

### Computational integration of multi-omics datasets

In the current study, using the Segway algorithm we demonstrated a use case of predicting tissue-specific super-enhancers based on multi-omics epigenomics measurements. The Segway genome segmentation method can also be applied to identify other types of regulatory elements if there is some prior knowledge to help assign semantics to the automatically inferred Segway labels. The multi-omics profiles, in conjunction with Segway, also have great potential to reveal disease-specific regulatory elements by identifying genomic labels that specifically occur in a group of patients vs. control samples, including drug toxicities.

Here, as a proof of concept, we also demonstrated that epigenomics and epitranscriptomics can be used as predictors for protein quantification. However, the Pearson correlation between predicted and observed protein level is still quite low. This is not entirely surprising, because many translational and post-translational effects are not yet captured, even with this rich, multi-omic analysis.

### Challenges

MultiomicsTracks96 is capable of generating high-dimensional multiome datasets. Still, one of the biggest challenges of multi-omics is the development of computational tools that can integrate such large-scale heterogenous datasets to define the flow of information and characterize multi-omic organization in health and disease, include the effects of drug treatments and environmental perturbations. There are significant advantages to preserving tissues as FFPE versus freezing, but for molecular analysis FFPE samples are more challenging. We have found excellent agreement between FFPE and frozen tissue for proteomic and DNA methylation measurements (R^2^ >0.9). Further improvements in efficiency of retrieval of soluble chromatin and RNA from FFPE samples are needed to facilitate integration of these heterogeneous datasets.

## CONCLUSIONS

Although it is well appreciated that in disease various omes are frequently perturbed, the molecular changes associated with the progression from health to disease are not sufficiently known to improve clinical outcomes. The decades-old practice of embedding and storing tissues in FFPE blocks generated vast biospecimen repositories that have been estimated to number in the hundreds of millions of samples (70). Long term storage (>10 years) does not appear to significantly affect our ability to extract macromolecules from FFPE blocks (26,52). In research settings, freezing is a common way to store biospecimens. As such, both modalities of tissue preservation and storing provide unprecedented biospecimen treasures for systems biology research. We have developed a 96-well format multi-omics toolbox, MultiomicsTracks96, for tissue sampling, analyte preparation, and analysis that generate multidimensional datasets. MultiomicsTracks96 makes it readily feasible to carry out high dimensional multiome analysis in large-scale clinical settings as well as animal model investigations that generate biospecimens, including FFPEs. While proteome is closer to cell phenotype than the nucleic acids, analysis reveals low correlation between protein levels and gene transcription profiles (73,74,76). Integration of additional epigenomic and epitranscriptomic targets to those used in this study should greatly improve understanding of protein expression profiles in organ development, health, environmental exposures, drug toxicity, disease and ageing.

## DECLARATIONS

### Ethics approval and consent to participate

All the animal care and experimental procedures were approved by University of Washington Institutional Animal Care and Use Committee (IACUC protocol is 4029-01) and carried out in compliance with the ARRIVE guidelines.

### Consent for publication

Not applicable

### Authors’ Contributions

KB conceived the project, designed the experiments, engineered the CryoGrid system, analyzed the data, and wrote the paper. DM designed, carried out experiments, contributed to CryoGrid design, analyzed the data, and wrote the paper. TV and IB designed the proteomic workflow and carried out the proteomic experiments. RZ and WSN carried out the Segway genome segmentation and protein expression prediction analysis. OD assisted in experimental design and edited the paper.

## Acknowledgments

We thank Dr. Mary Regier, UW ISCRM Genomics Core, for assistance with bioinformatics. We thank the Altemeier Lab UW Medicine SLU for providing mouse organs,

## Inclusion and Diversity

One or more of the authors of this paper self-identifies as an underrepresented ethnic minority in science. One or more of the authors of this paper self-identifies as a member of the LGBTQ+ community. One or more of the authors of this paper self-identifies as living with a disability. One or more of the authors of this paper received support from a program designed to increase minority representation in science.

## SUPPLEMENT

**Fig.S1.**
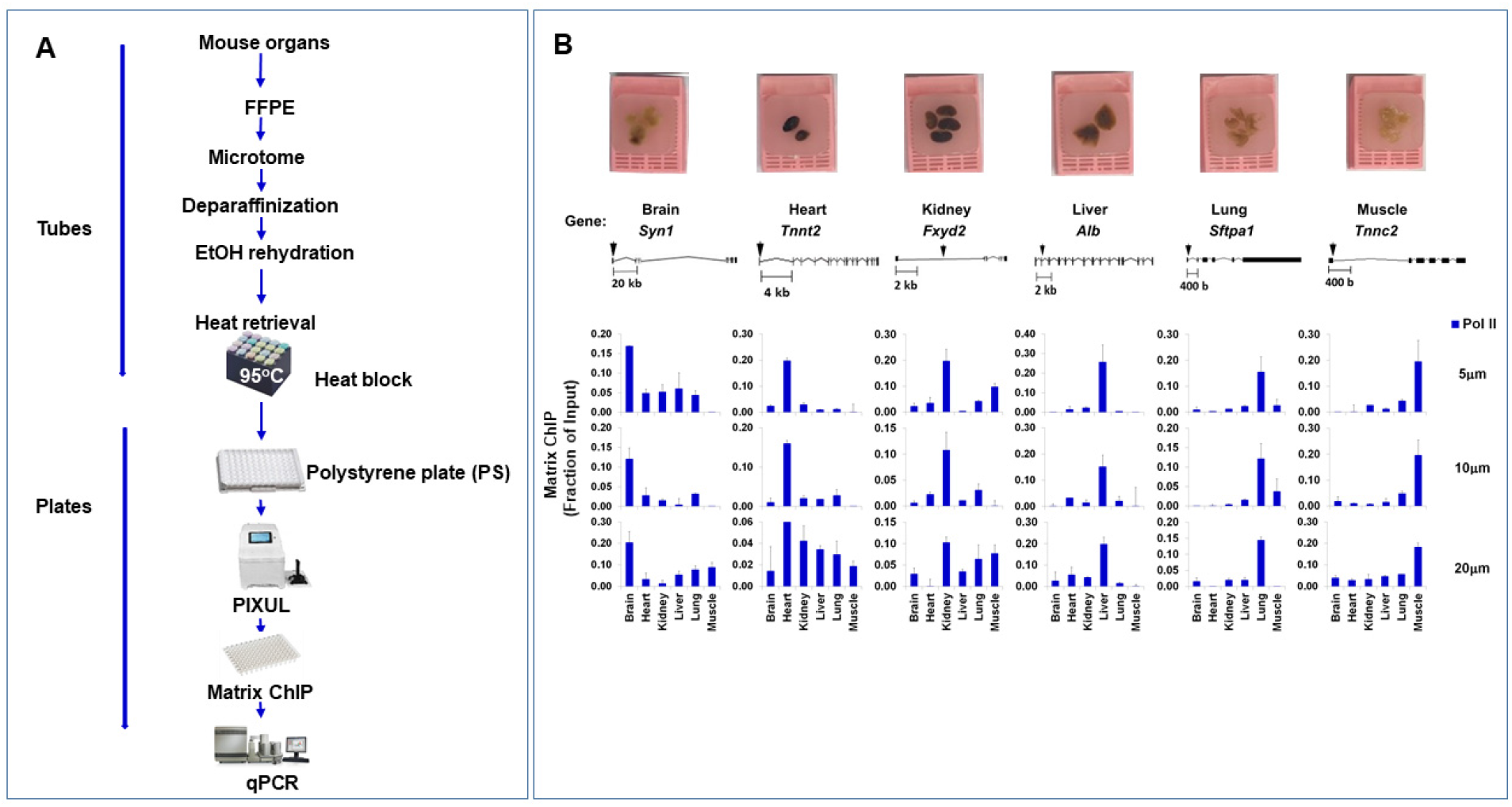
PIXUL-FFPE-ChIP-qPCR analysis at mouse organs specific genes: Microtome sections thickness. ***A.*** FFPE blocks of mouse brain, heart, kidney, liver, lung, and muscle were cut with a microtome at 5, 10, and 20μm section thickness, total of 8 curls. After deparaffinization with Xylene, EtOH rehydration, and heat retrieval, samples were treated in PIXUL to isolate soluble chromatin, which was used in Matrix ChIP using anti-RNA Polymerase II CTD (Pol II CTD) antibody. ***B.*** FFPE blocks. Cartoons of the genes. Black arrows show position of qPCR primers used in ChIP. Inputs were diluted 20X to overcome PCR interference. Results (expressed as a fraction of input) represent mean+SEM (n=2).

**Fig.S2.**
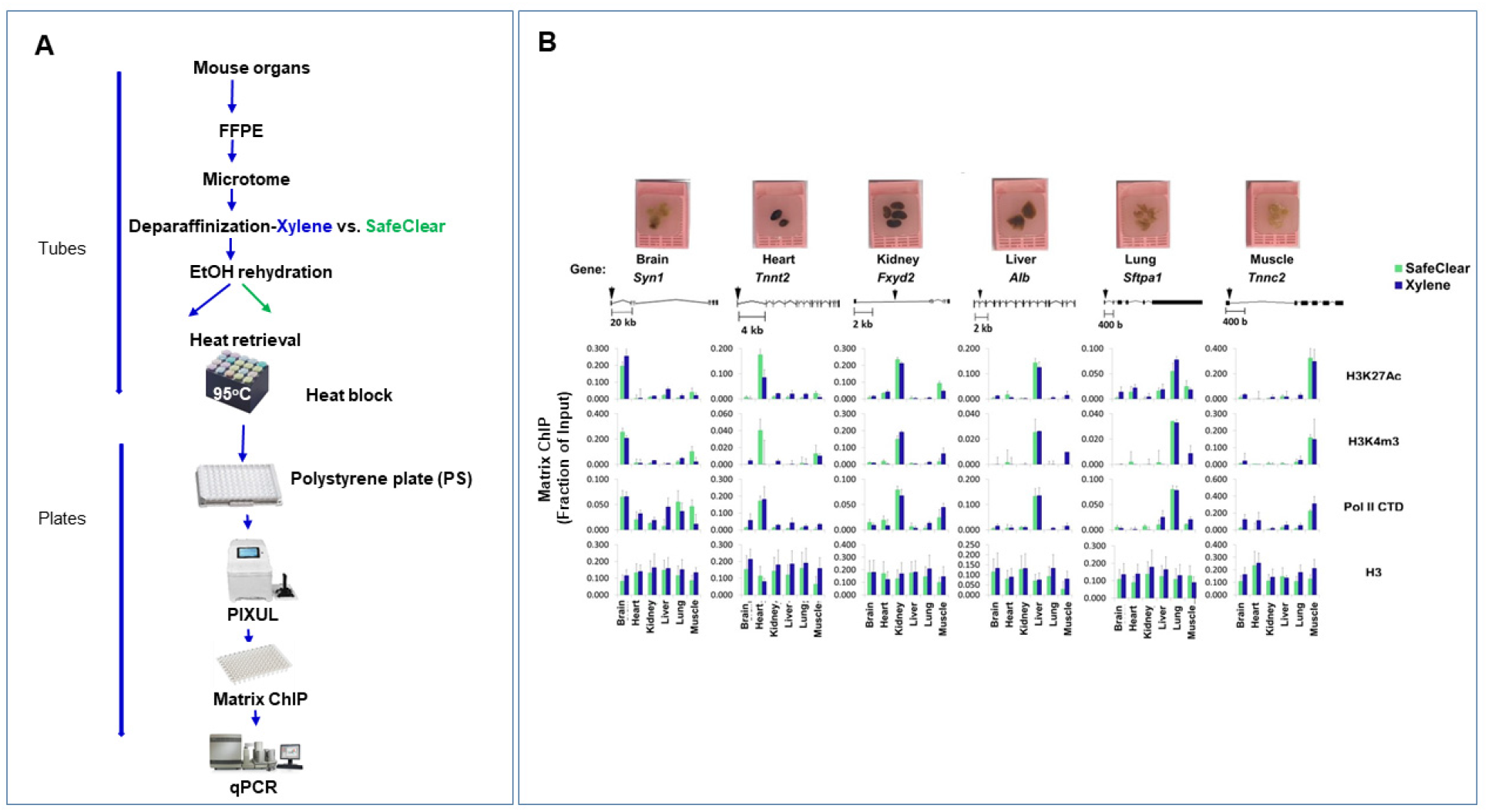
PIXUL-FFPE-ChIP analysis at mouse organs specific genes: SafeClear compared to Xylene. ***A.*** FFPE slices (5μm) were generated from FFPE blocks using a microtome, total 2 series for each organ. After deparaffinization with either SafeClear or Xylene, EtOH rehydration, and heat retrieval, samples were PIXULed. ***B.*** Extracted chromatin was analyzed in Matrix ChIP using antibodies to H3K27Ac, H3K4m3, Pol II CTD, and histone H3. Mouse IgG was used for background subtraction. Inputs were diluted 20X to overcome PCR interference. Results (expressed as a fraction of input) represent mean+SEM (n=2 different FFPE extractions).

**Fig.S3.**
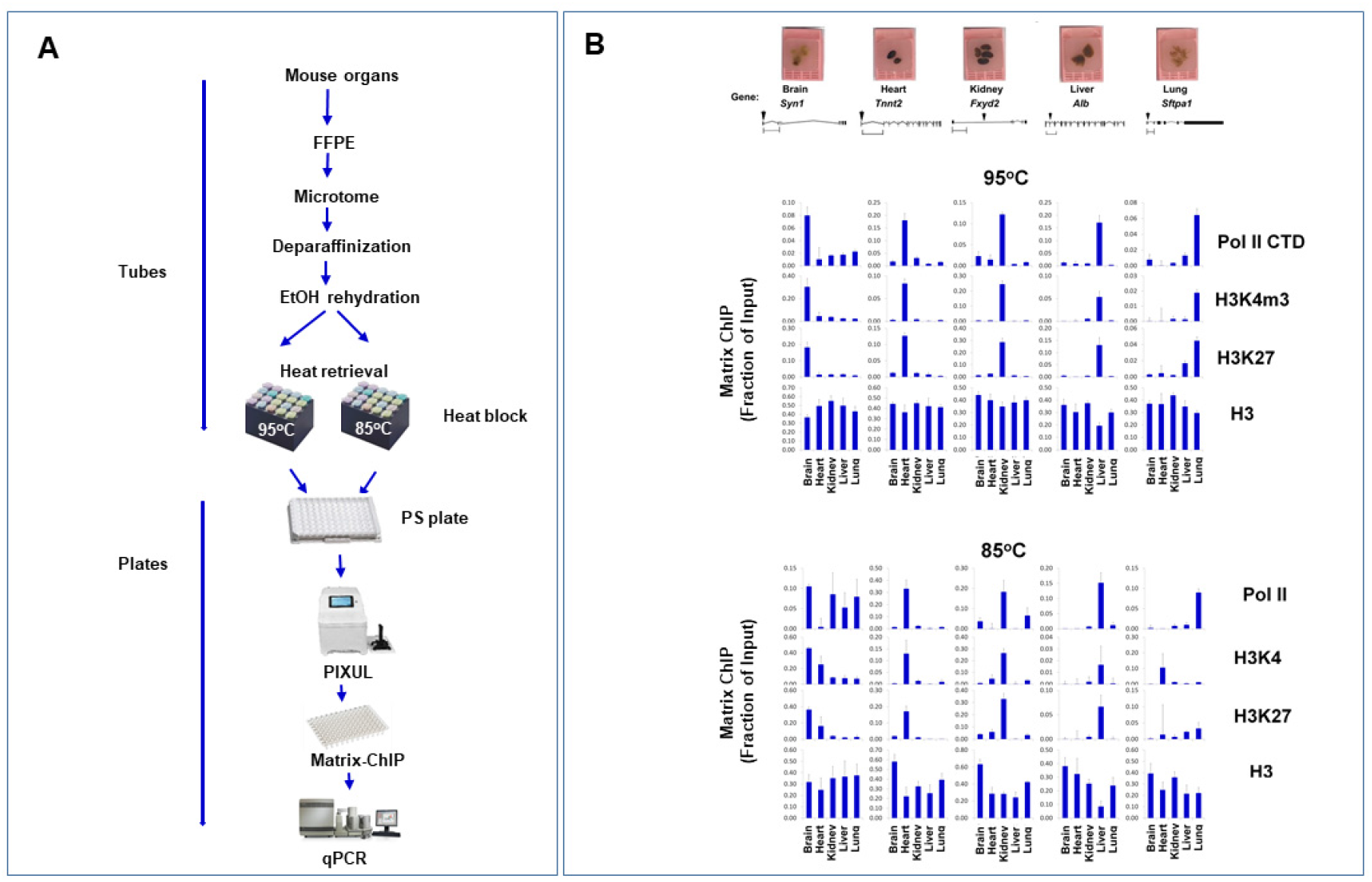
PIXUL-FFPE-ChIP analysis at mouse organs specific genes: Chromatin heat retrieval temperature. ***A.*** FFPE slices (5μm) were generated from FFPE blocks using a microtome, total 2 series for each organ. After deparaffinization with SafeClear and EtOH rehydration, heat retrieval was compared at 95°C vs. 85°C and then samples were PIXULed. ***B.*** Extracted soluble chromatin was analyzed in Matrix ChIP using antibodies to Pol II CTD, H3K4m3, H3K27Ac, and histone H3. Mouse IgG was used for background subtraction. Inputs were diluted 20X to overcome PCR interference. Results (expressed as a fraction of input) represent mean+SEM (n=4 qPCRs).

**Fig.S4.**
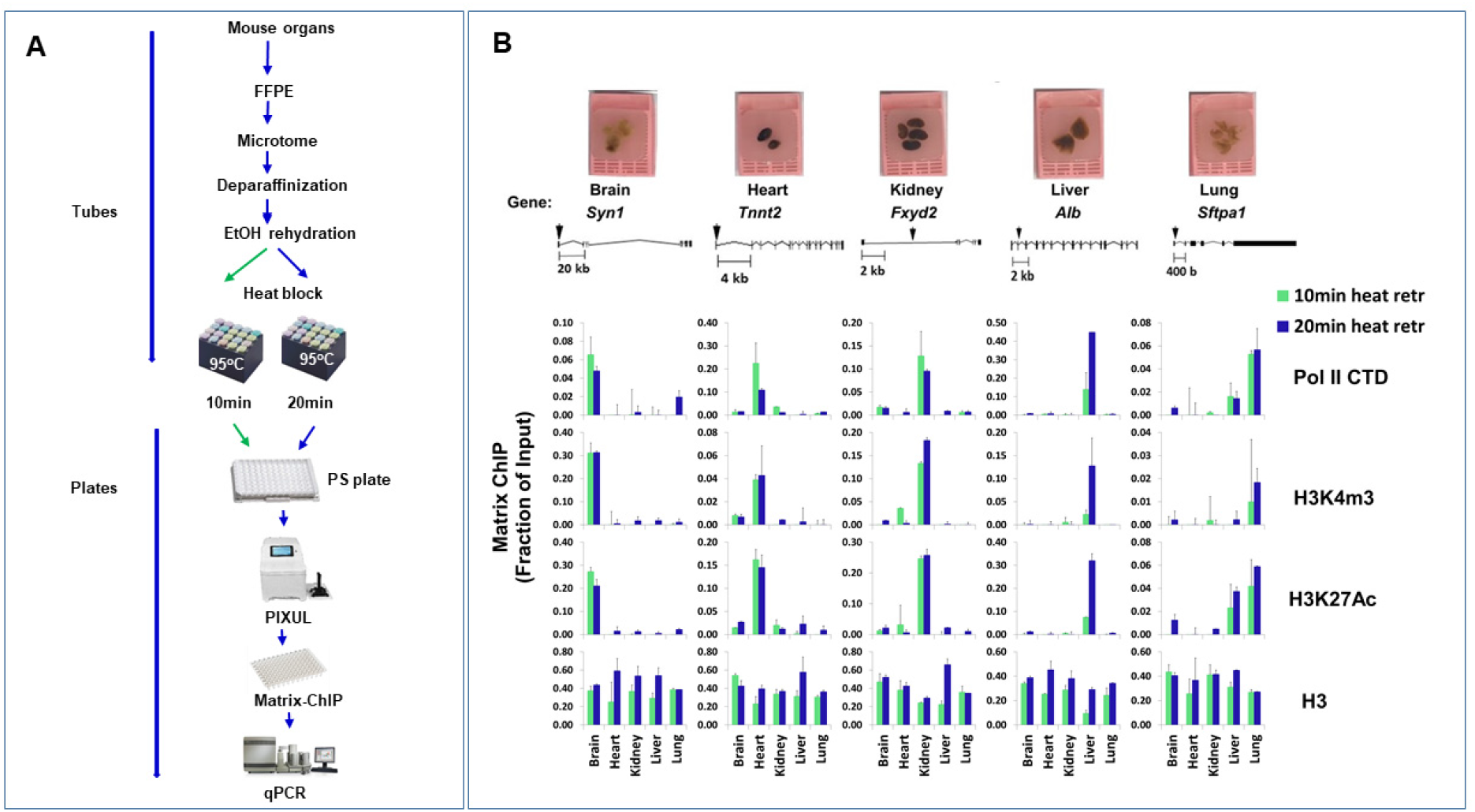
PIXUL-FFPE-ChIP analysis at mouse organs specific genes: Chromatin heat retrieval time. ***A.*** FFPE slices (5μm) were generated from FFPE blocks using a microtome, total 2 series for each organ. After deparaffinization with SafeClear and EtOH rehydration, heat retrieval was compared at 10 vs. 20min (95°C) and then samples were PIXULed. ***B.*** Extracted soluble chromatin was analyzed in Matrix ChIP using antibodies to Pol II CTD, H3K4m3, H3K27Ac, and histone H3. Mouse IgG was used for background subtraction. Inputs were diluted 20X to overcome PCR interference. Results (expressed as a fraction of input) represent mean+SEM (n=4 qPCRs).

**Fig.S5.**
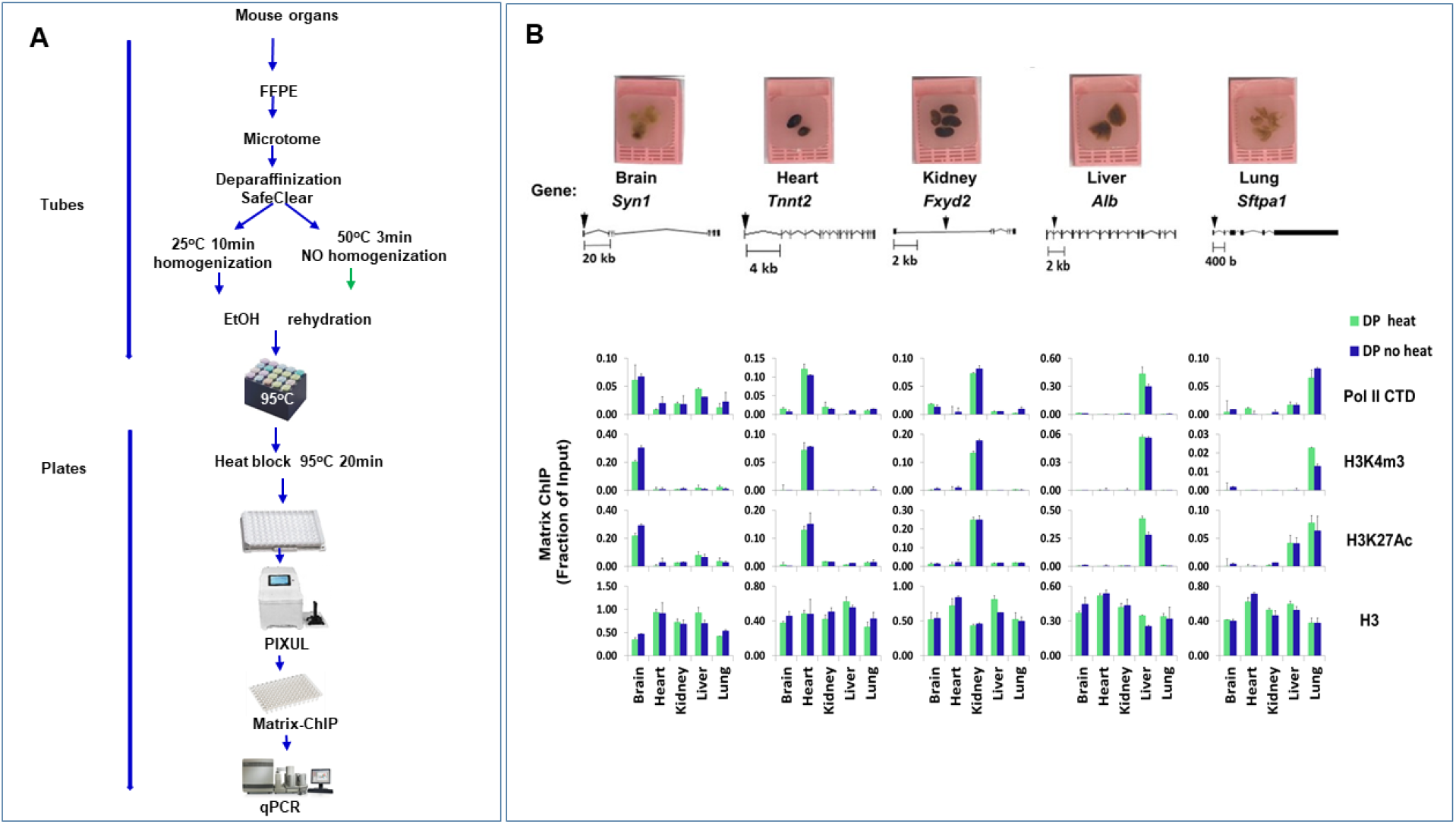
PIXUL-FFPE-ChIP analysis at mouse organs specific genes: Deparaffinization temperature. ***A.*** FFPE slices (5μm) were generated from FFPE blocks using a microtome, total 2 series for each organ. Deparaffinization with SafeClear was compared at 25°C (10min) vs. 50°C (3min) without manual homogenization. After EtOH rehydration and heat retrieval (95°C for 20min), samples were PIXULed as before. ***B.*** Extracted soluble chromatin was analyzed in Matrix ChIP using antibodies to Pol II CTD, H3K4m3, H3K27Ac, and histone H3. Mouse IgG was used for background subtraction. Inputs were diluted 20X to overcome PCR interference. Results (expressed as a fraction of input) represent mean+SEM (n=4 qPCRs).

**Fig.S6.**
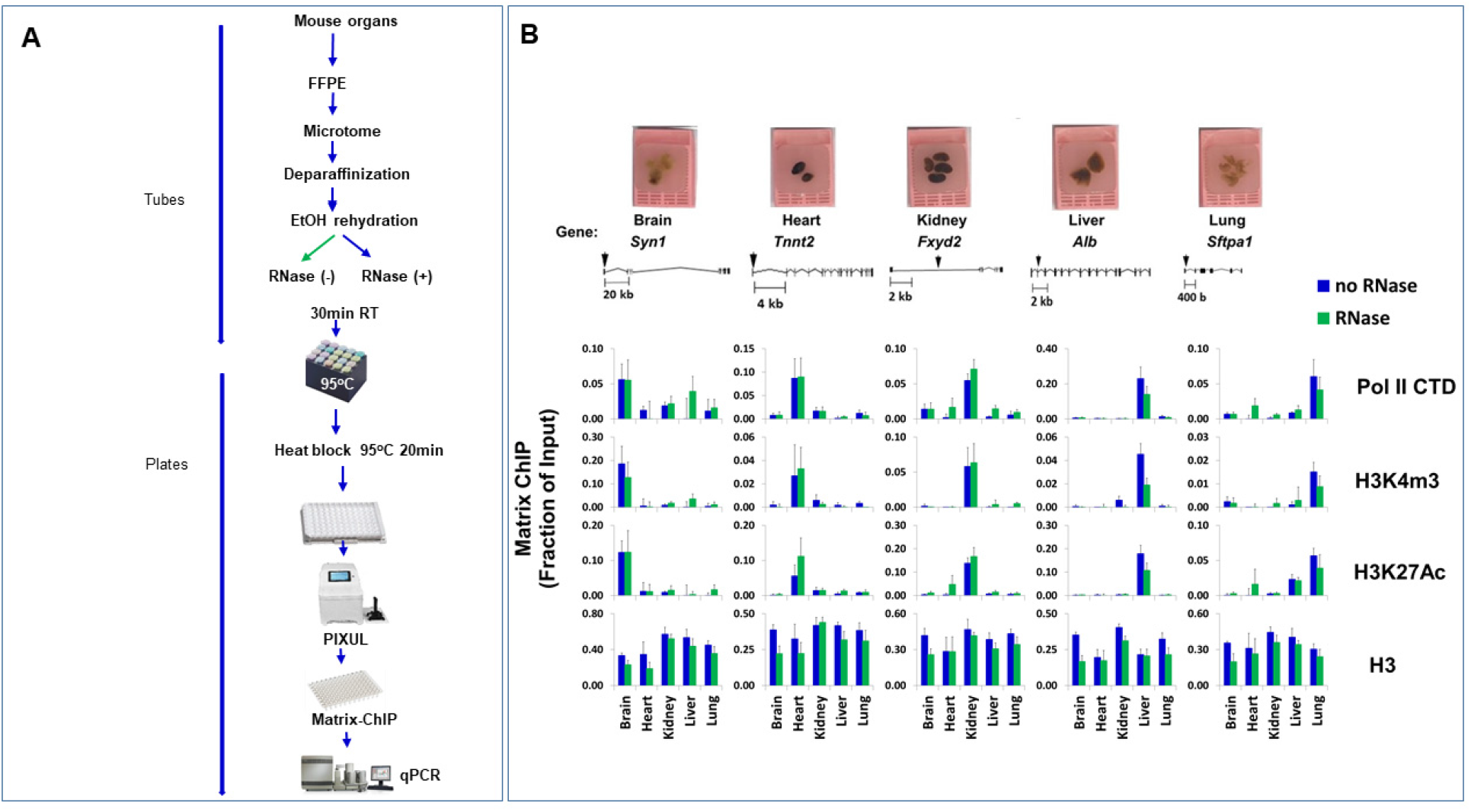
PIXUL-FFPE-ChIP analysis at mouse organs specific genes: RNase treatment. ***A.*** FFPE slices (5μm) were generated from FFPE blocks using a microtome, total 2 series for each organ. After deparaffinization with SafeClear and EtOH, rehydrated samples were treated with or without RNase (30min, RT). Retrieval was done at 95°C for 20min and then samples were PIXULed. ***B.*** Extracted soluble chromatin was analyzed in Matrix ChIP using antibodies to Pol II CTD, H3K4m3, H3K27Ac, H3K4m3, and histone H3. Mouse IgG was used for background subtraction. Inputs were diluted 20X to overcome PCR interference. Results (expressed as a fraction of input) represent mean+SEM (n=4 qPCRs).

**Fig.S7.**
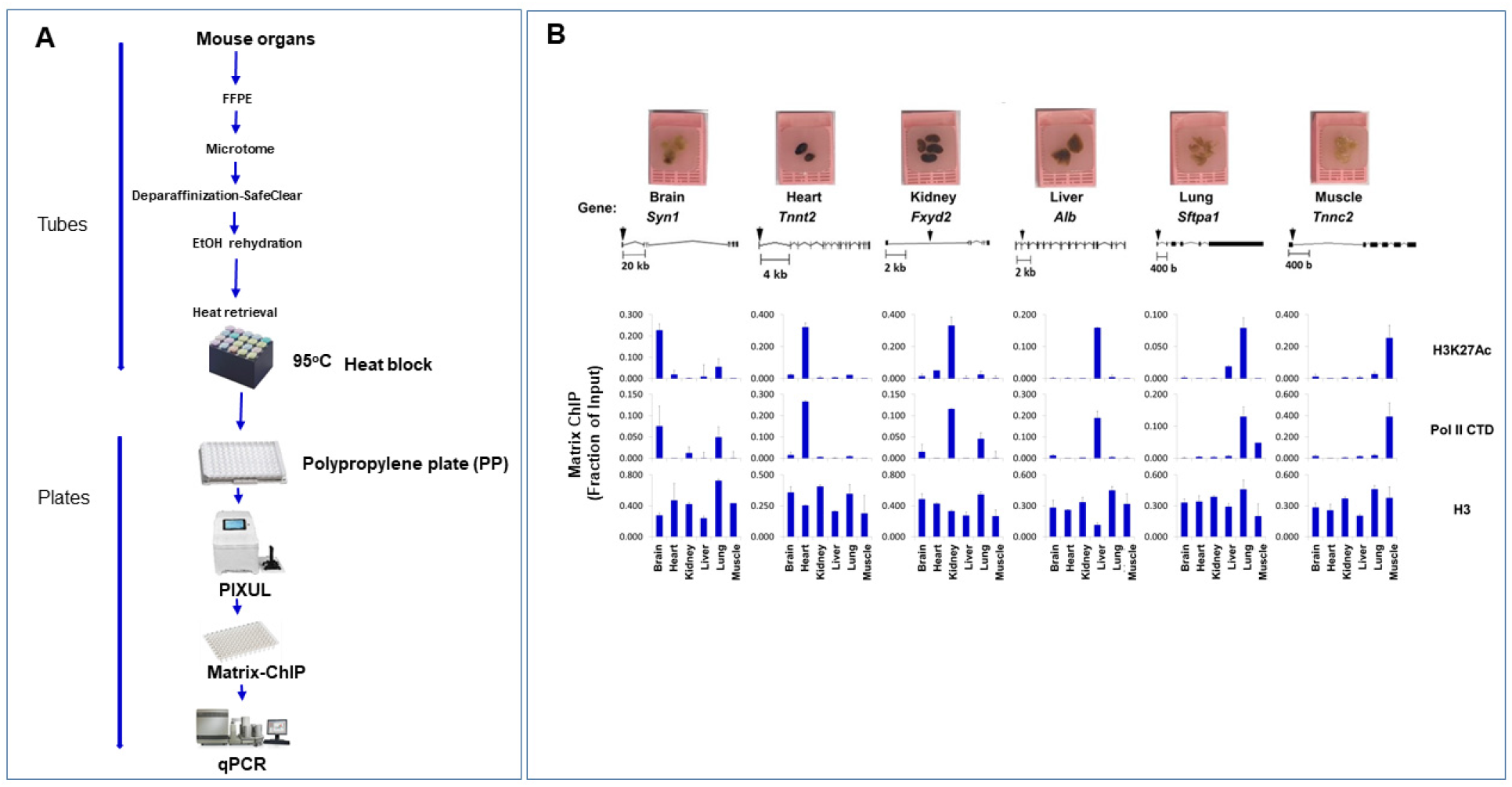
PIXUL-FFPE-ChIP analysis at mouse organs specific genes: Polypropylene PIXUL plates. ***A.*** FFPE slices (5μm) were generated from FFPE blocks using a microtome, total 2 series for each organ. After deparaffinization with SafeClear, EtOH rehydration, and heat retrieval in a heat block (95°C for 20min), samples were PIXULed using 96 well polypropylene plate. ***B.*** Extracted soluble chromatin was analyzed in Matrix ChIP using antibodies to H3K27Ac, Pol II CTD, and histone H3. Mouse IgG was used for background subtraction. Inputs were diluted 20X to overcome PCR interference. Results (expressed as a fraction of input) represent mean+SEM (n=4 qPCRs).

**Fig.S8.**
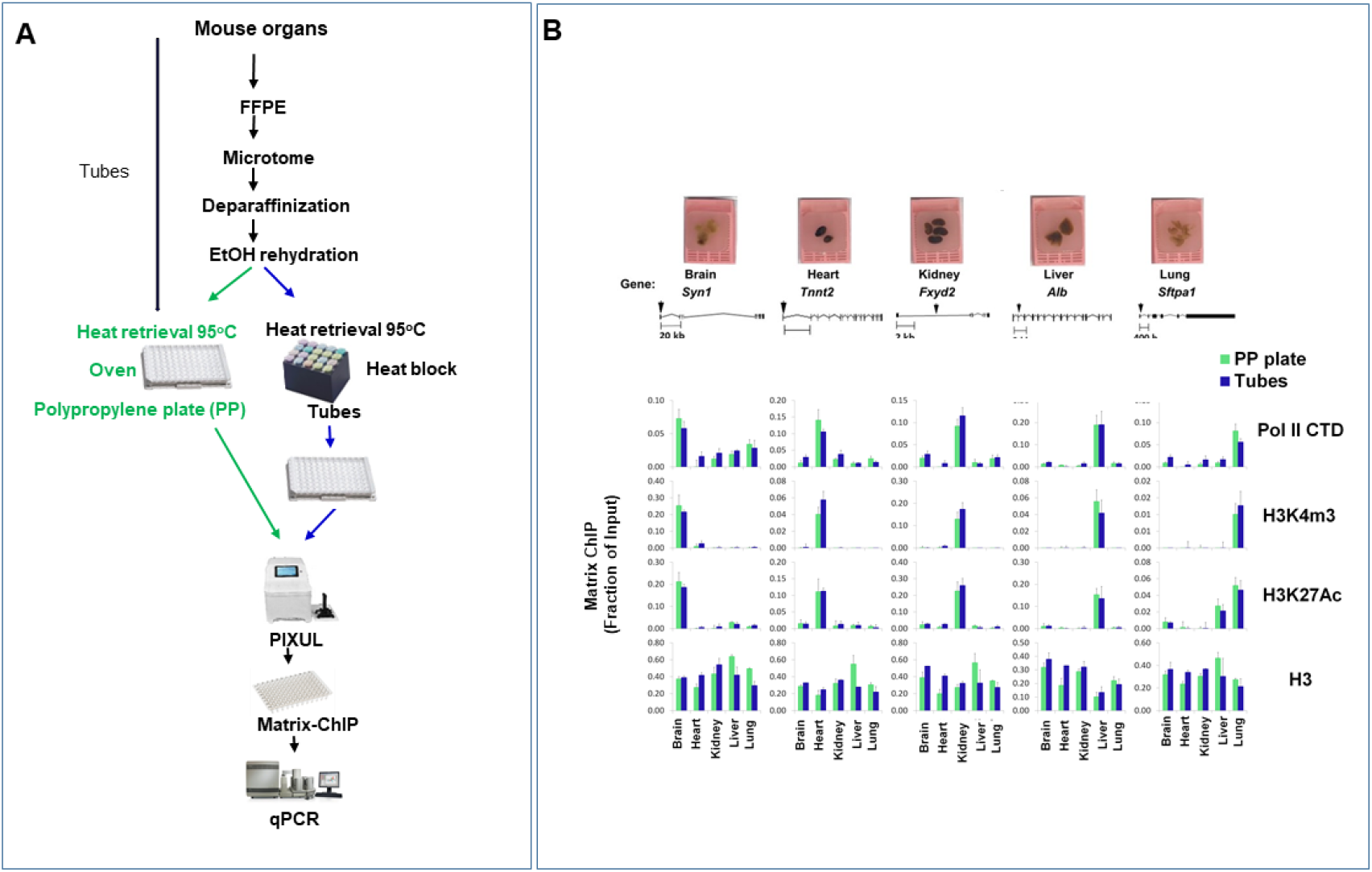
PIXUL-FFPE-ChIP analysis at mouse organs specific genes: Heat retrieval using heat-resistant polypropylene PIXUL plates. ***A.*** FFPE slices (5μm) were generated from FFPE blocks using a microtome, total 2 series for each organ. After deparaffinization with SafeClear and EtOH rehydration, heat retrieval was compared using heat resistant polypropylene plate at 95°C (20 min) in an oven compared to test tubes in a heat-block (95°C for 20min). The test tube samples were transferred to a 96 well polypropylene plate for PIXUL while the heated plate was directly PIXULed after cooling. ***B.*** Extracted soluble chromatin was analyzed in Matrix well plate for PIXUL-ChIP using antibodies to Pol II CTD, H3K4m3, H3K27Ac, and histone H3. Mouse IgG was used for background subtraction. Inputs were diluted 20X to overcome PCR interference. Results (expressed as a fraction of input) represent mean+SEM (n=4 qPCRs).

**Fig.S9.**
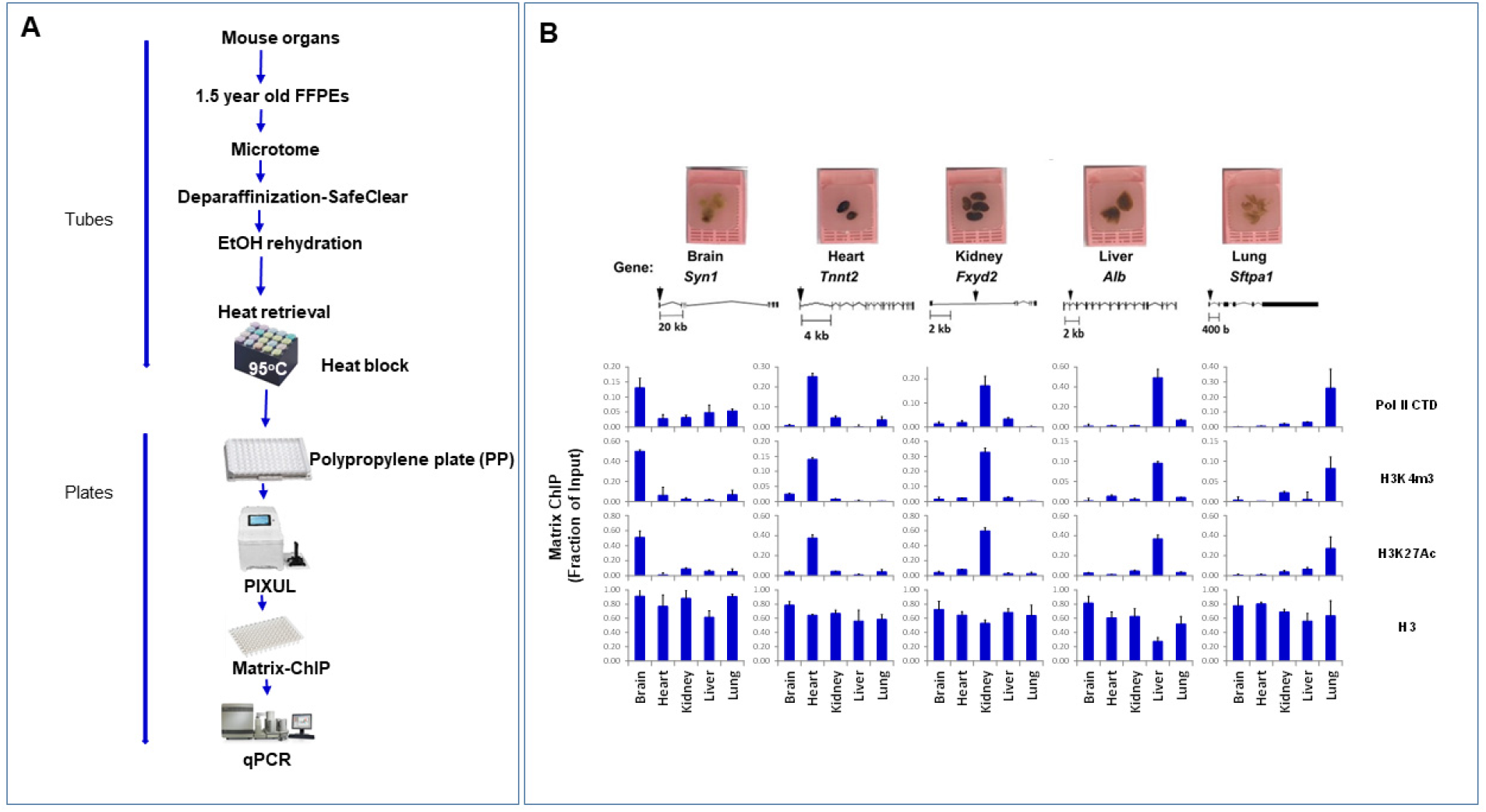
PIXUL-FFPE-ChIP analysis at mouse organs specific genes: 1.5-year-old FFPEs. ***A.*** FFPE slices (5μm) were generated from 1.5-year-old FFPE blocks using a microtome, total 2 series for each organ. After deparaffinization with SafeClear, EtOH rehydration, and heat retrieval in a heat block (95°C for 20min), samples were PIXULed using 96 well polypropylene plate. ***B.*** Extracted soluble chromatin was analyzed in Matrix ChIP using antibodies to Pol II CTD, H3K4m3, H3K27Ac, and histone H3. Mouse IgG was used for background subtraction. Inputs were diluted 20X to overcome PCR interference. Results (expressed as a fraction of input) represent mean+SEM (n=4 qPCRs).

**Fig.S10.**
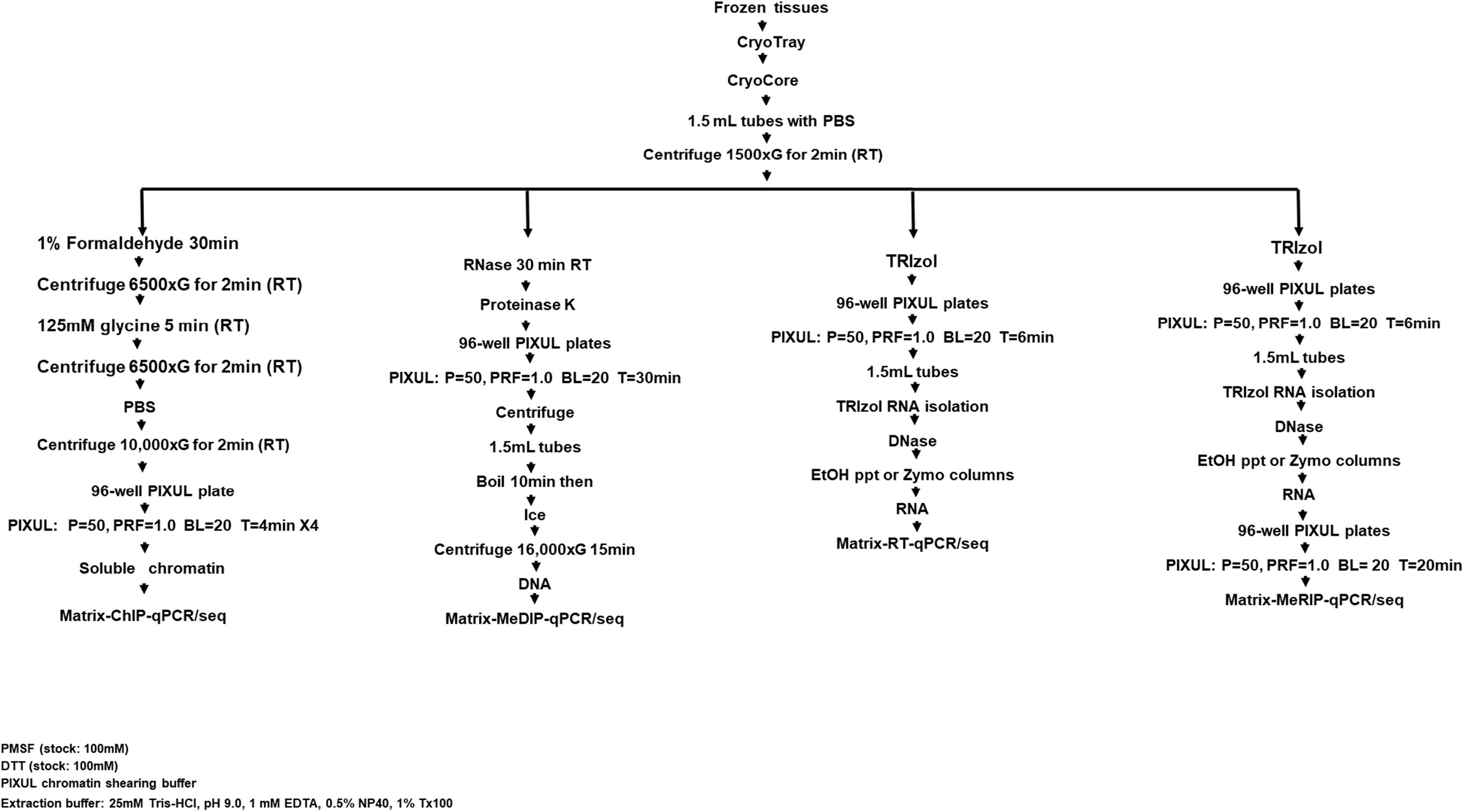
Multi-omics workflows for frozen tissues.

**Fig.S11.**
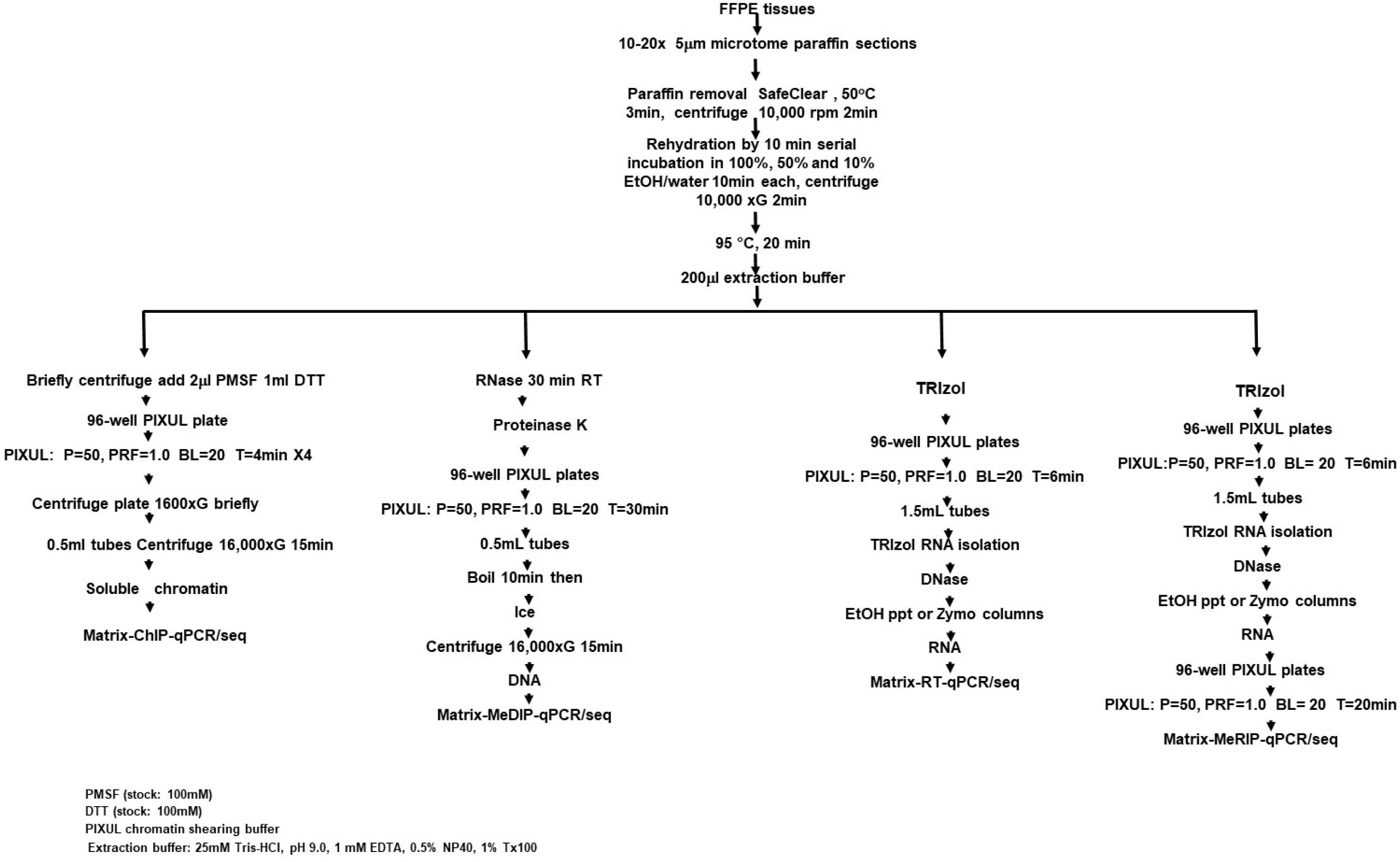
Multi-omics workflows for FFPE tissues.

**Fig.S12.**
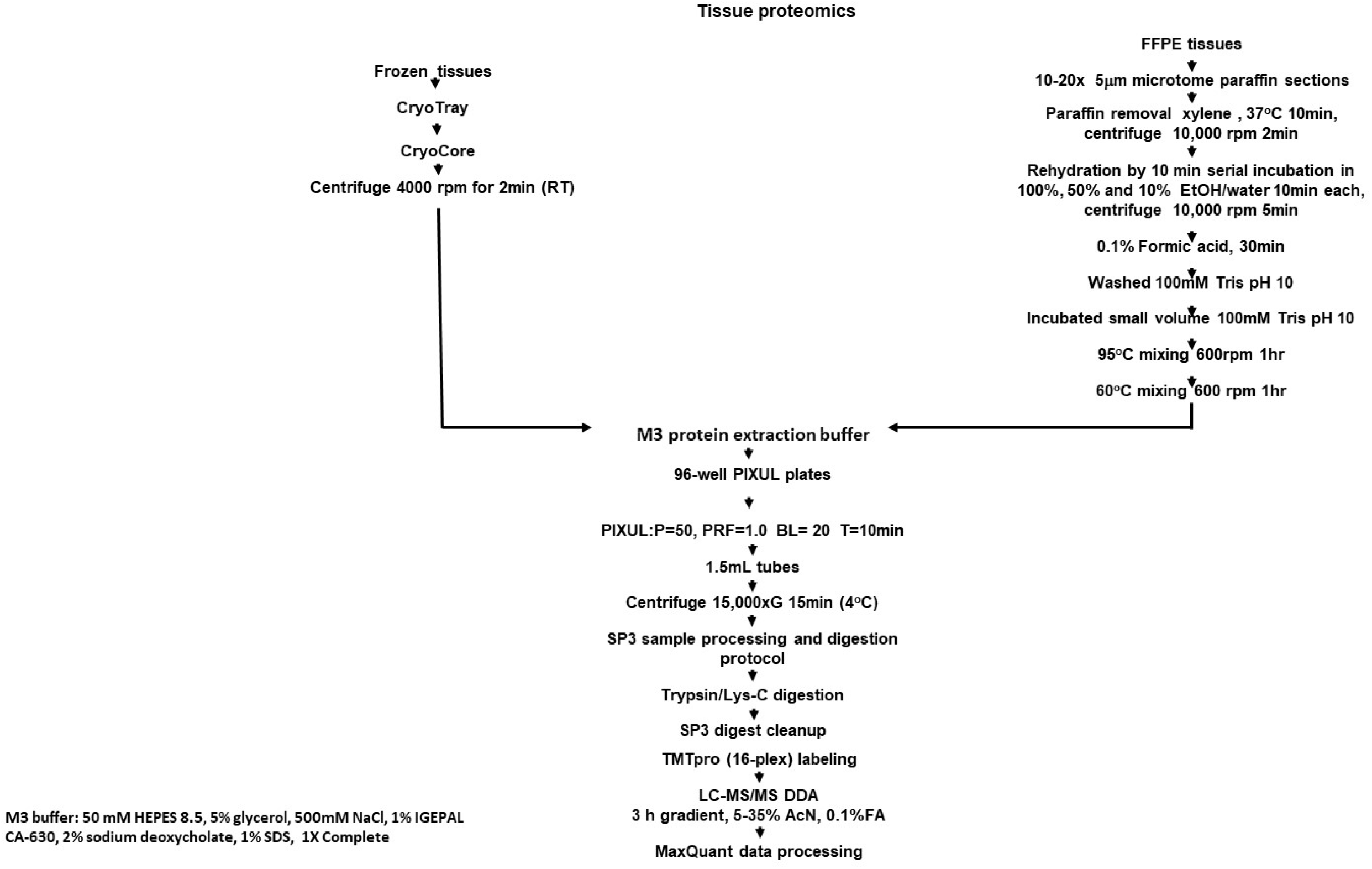
Proteomics workflows for frozen and FFPE tissues.

**Fig.S13.**
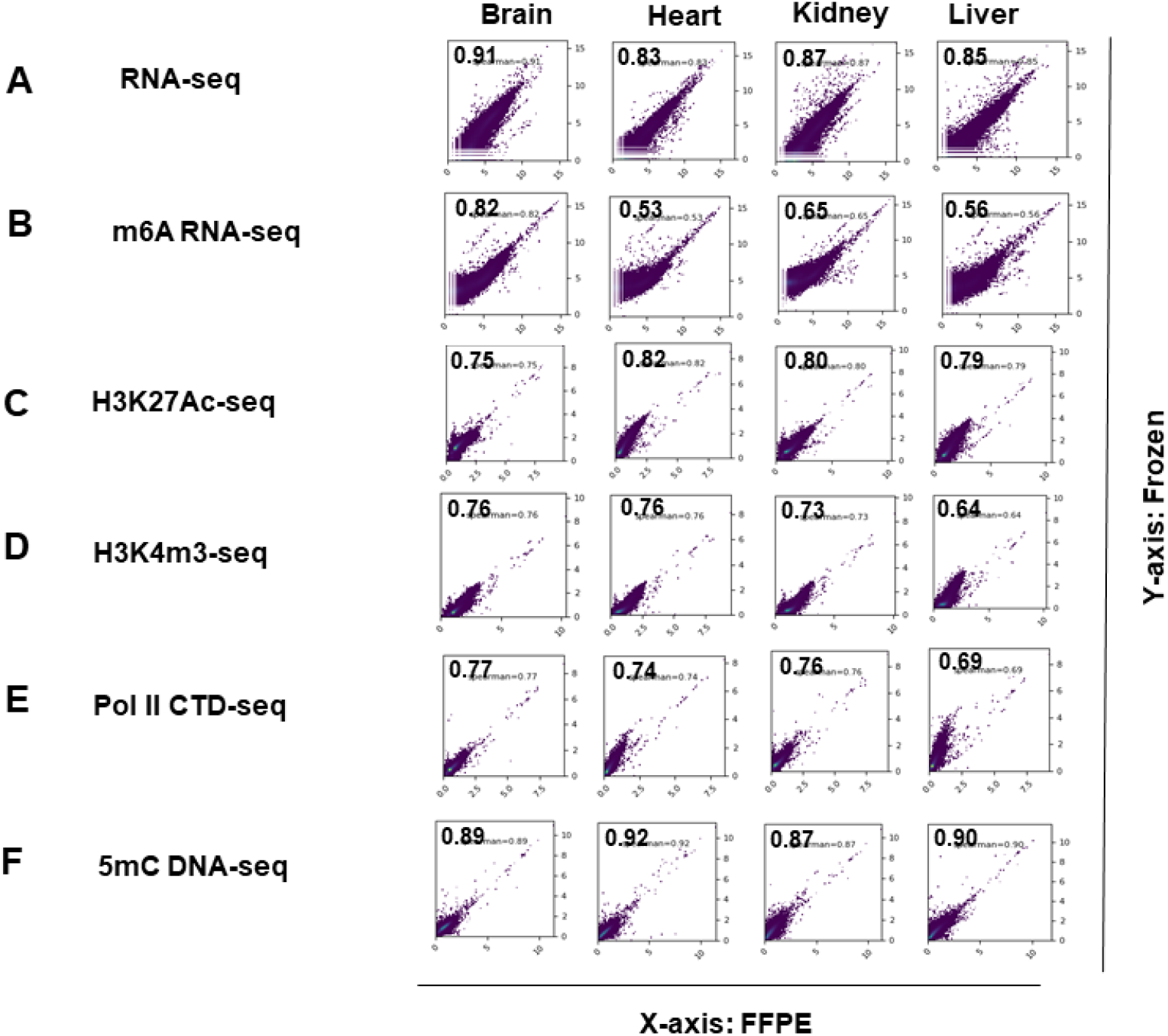
Scatter plots of comparative analysis of PIXUL-Matrix-seq generated 6-omics nucleic acids datasets from mouse organs frozen and FFPE blocks. ***A.*** RNA-seq. ***B.*** m6A RNA-seq. ***C.*** H3K27Ac. ***D.*** H3K4m3-seq. ***E.*** Pol II CTD-seq. ***F.*** 5mC DNA-seq. Numbers in the plots field represent Spearman coefficients.

**Fig.S14. Multi-omics.**
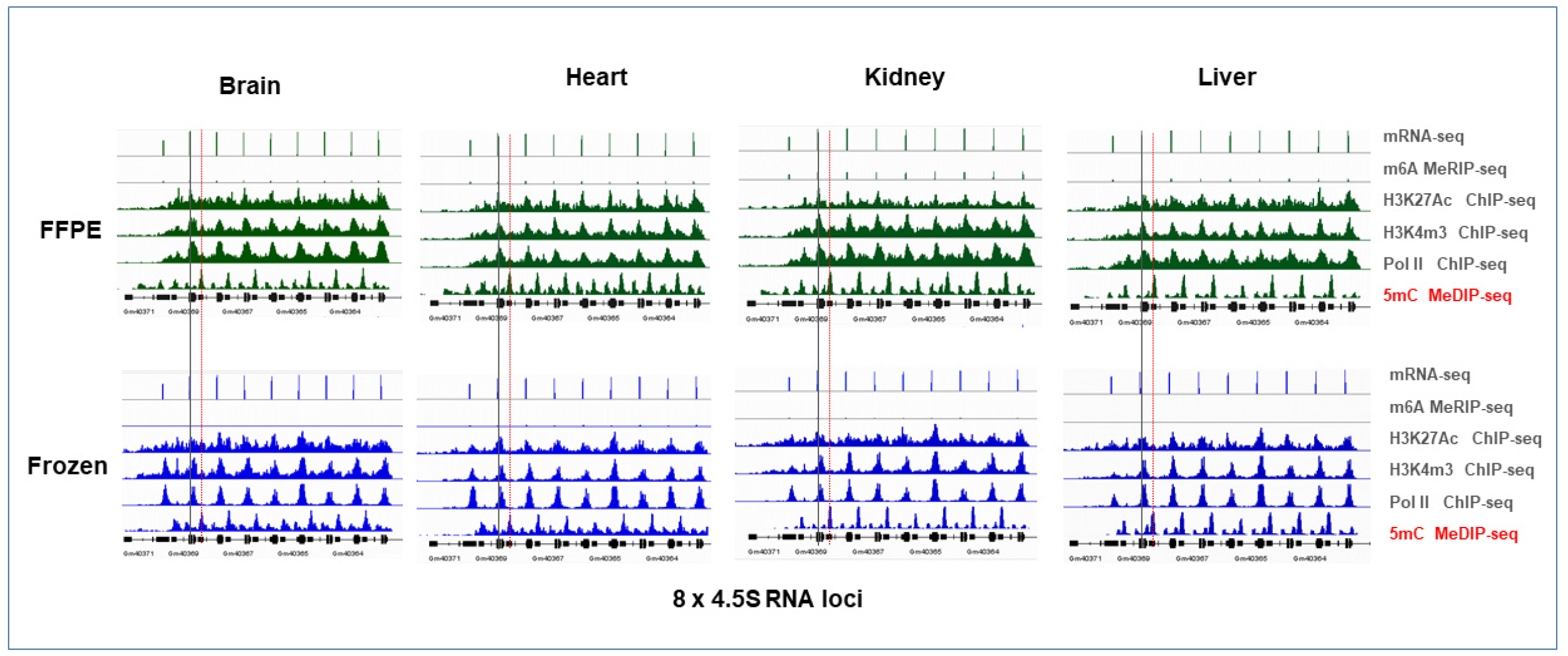
PIXUL-Matrix-seq. PIXUL was used to prepare RNA, DNA, and chromatin samples from mouse fresh frozen and FFPE brain, heart, kidney, and liver tissues. Matrix was used for RT (mRNA-seq) and immunoprecipitation of methylated RNA (MeRIP: m6A), methylated DNA (MeDIP: 5mC), and chromatin immunoprecipitation (ChIP: H3K27Ac, H3K4m3, and Pol II CTD). Libraries were generated and sequenced (NextSeq2000). IGV snapshot is shown at the ubiquitously expressed 4.5 ribosomal RNA (4.5 S rRNA) loci in each mouse organ. 4.5S rRNAs are well expressed in each organ. m6A-RNA-seq shows that 4.5S rRNA is not m6A methylated. The transcription (Pol II) of 4.5S rRNA loci is associated with permissive epigenetic marks (H3K27Ac and H3K4m3). At the 4.5S rRNA loci RNA, Pol II and the permissive marks line up (*black*) but 5mC (*red*) does not, suggesting epigenetic antagonism.

**Fig.S15.**
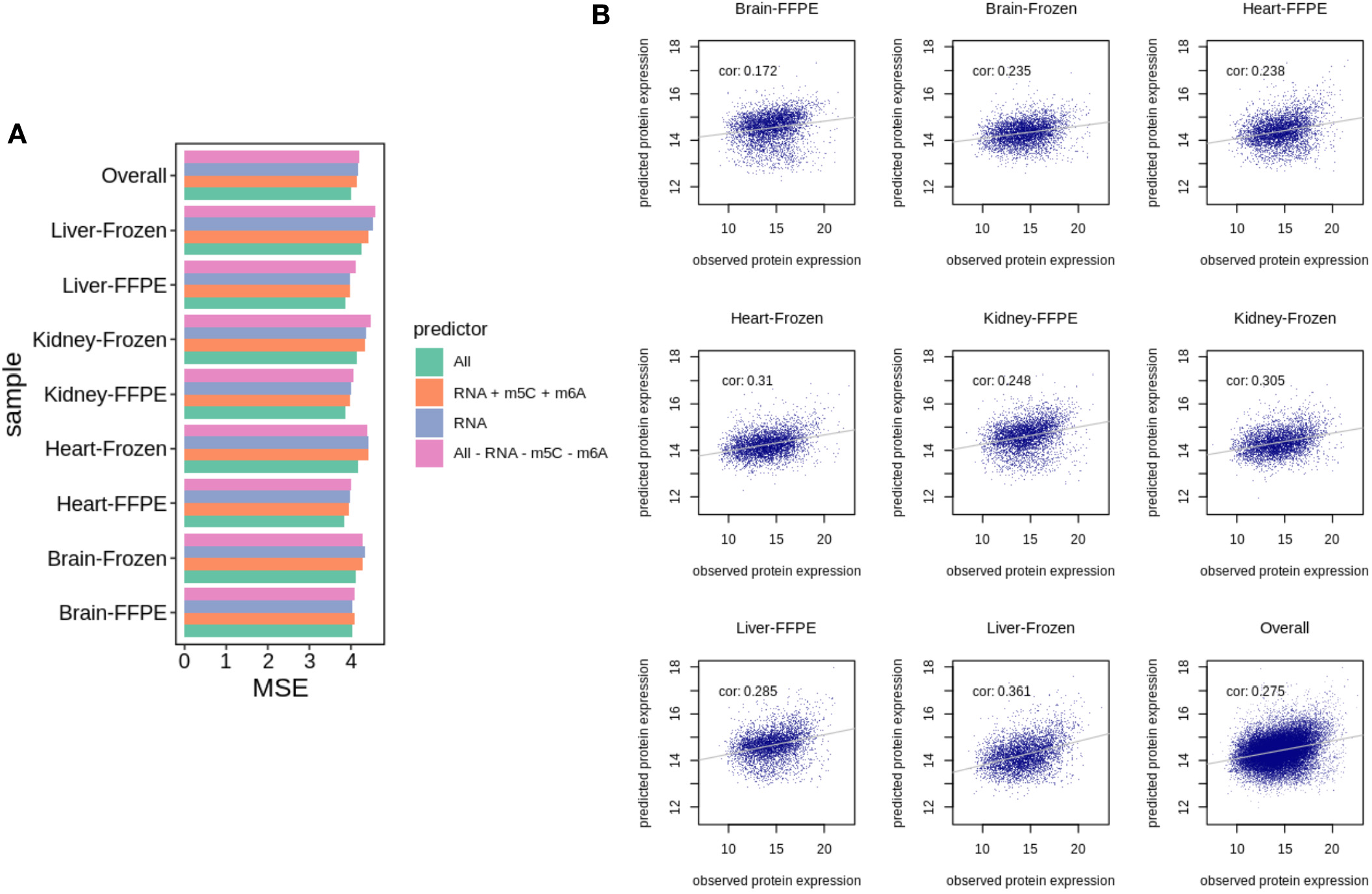
Protein expression prediction performance. ***A.*** mean-squared-error (MSE) of predicted vs. observed protein expression profile, separated by organ, sample prep, and different sets of predictors***. B.*** scatterplot of predicted vs. observed protein expression per organ and sample prep; predicted protein expression is based on all other seven multi-omics assays (corresponding to “All” predictor). The grey line on each plot indicates the trend using a linear regression model. Pearson correlation coefficients are calculated across all proteins within and across organ and sample prep.

**Fig.S16.**
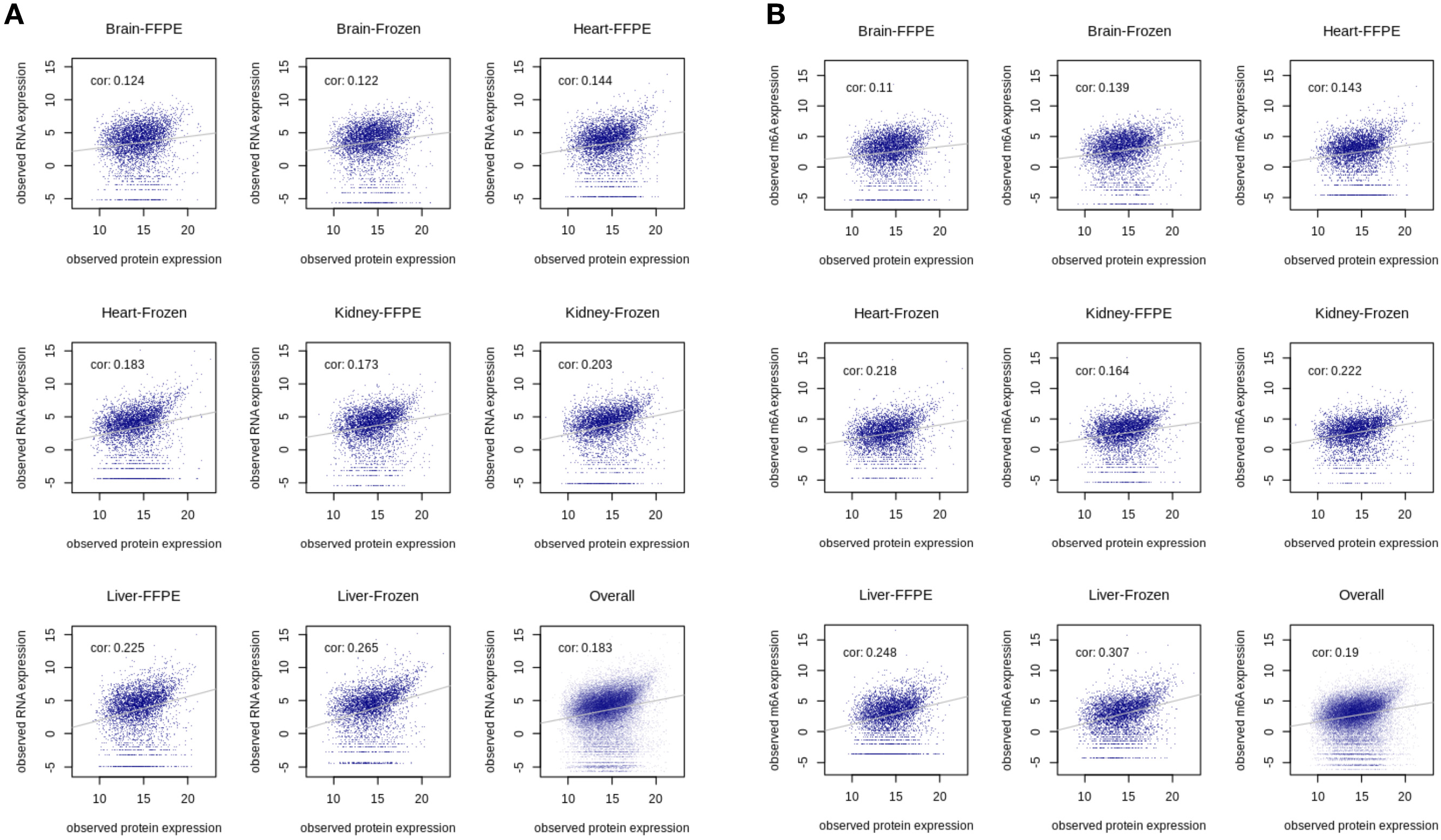
Scatter plot between observed protein expression vs. observed mRNA expression (A) and observed m6A expression (B). The grey line on each plot indicates the trend using a linear regression model. Pearson correlation coefficients are calculated across all proteins within and across organ and sample prep.

## TABLES

**Table S1.** Hardware and labware.

| Item Description | Manufacturer/Supplier | Location | Part No. |
| --- | --- | --- | --- |
| PIXUL™ 96-well sonicator | Matchstick Technologies | Kirkland, WA | P01199-01A |
|  | Active Motif | Carlsbad, CA | 53130 |
| CryoTray | Matchstick Technologies | Kirkland, WA | P01200-01A |
| CryoBlock | Matchstick Technologies | Kirkland, WA | P01201-01A |
| CryoCore | Matchstick Technologies | Kirkland, WA | P01202-01A |
| CryoCore trephines | Matchstick Technologies | Kirkland, WA | P01203-01A |
| PlateHandle | Matchstick Technologies | Kirkland, WA | P01204-01A |
| Fluke 52-II Dual Probe Thermometer | Fluke Corporation | Everett, WA | 674689 |
| Fluke 80PJ-1 Bead Probe | Fluke Corporation | Everett, WA | 750422 |
| Small Dual Angle Beta Radiation Shield (used as iPad stand) | Universal Medical Inc. | Oldsmar, FL | UM3400 |
| iPAD Pro | Apple | Cupertino, CA | Model A2377 |
| NextSeq2000 | Illumina | San Diego, CA | 20038897 |
| 4200 TapeStation | Agilent | Santa Clara, CA | G29991BA |
| Invitrogen™ Qubit™ 3 Fluorometer | ThermoFisher | Waltham, MA | Q33216 |
| NanoDrop 1000 Spectrophotometer | ThermoFisher | Waltham, MA | E112352 |
| Eppendorf ThermoMixer® C | Eppendorf | Hamburg, Germany | 5382000023 |
| Costar 96-well polypropylene round-bottom (used in PIXUL) | Corning | Oneonta, NY | 3365 |
| GeneMate 96-well 0.2ml plate semi skirt | VWR | Radnor, PA | 490003-794 |
| MicroAmp Optical Adhesive Film (used in PIXUL) | Applied Biosystems | Carlsbad, CA | 4311971 |
| E1-ClipTip™ Electronic Multichannel Pipettes | ThermoFisher | Waltham, MA | 4671090 |
| 7900HT Fast Real-Time PCR System with 384-Well Block Module | ThermoFisher | Waltham, MA | 4329001 |
| Oasis uElute SPE | Waters | Milford, MA | 186008052 |
| EasyLC 1200 | Thermo Scientific | San Jose, CA | LC140 |
| Orbitrap Exploris 480 | Thermo Scientific | San Jose, CA | BRE725539 |

**Table S2.** Kits and enzymes.

| Item Description | Manufacturer/Supplier | Location | Part No. |
| --- | --- | --- | --- |
| Zymo-Seq RiboFree Total RNA Library Kit | Zymo Research | Tustin, CA | R3003 |
| Zymo RNA Clean & Concentrator-5 | Zymo Research | Tustin, CA | R1016 |
| PureLink™ RNA Micro Kit | Invitrogen | Carlsbad, CA | 12183016 |
| SpeedBeads™ magnetic carboxylate modified particles | MilliporeSigma | Burlington, MA | GE65152105050250 |
| SuperScript IV Reverse Transcriptase | ThermoFisher | Waltham, MA | 18090050 |
| RNaseOUT™ Recombinant Ribonuclease Inhibitor | ThermoFisher | Waltham, MA | 10777-019 |
| RNase-Free DNase I | Lucigen | Middleton, WI | E0013-1D4 |
| DNase I 10X Reaction Buffer | Lucigen | Middleton, WI | SS000751-D2 |
| Invitrogen™ Proteinase K, recombinant-Invitrogen | ThermoFisher | Waltham, MA | 25530015 |
| Standard Sensitivity Large Fragment Analysis Kit | Agilent | Santa Clara, CA | DNF-492 |
| High Sensitivity RNA ScreenTape Ladder | Agilent | Santa Clara, CA | 5067-5581 |
| High Sensitivity RNA ScreenTape Sample Buffer | Agilent | Santa Clara, CA | 5067-5580 |
| High Sensitivity RNA ScreenTape | Agilent | Santa Clara, CA | 5067-5579 |
| Next Gen DNA Library Kit | Active Motif | Carlsbad, CA | 53216 |
| xGen ssDNA Low-Input DNA Lib Prep 16rxn, | IDT | Coralville, IA | 10009859 |
| xGen UDI Primers, 16 rxn | IDT | Coralville, IA | 10005975 |
| NextSeq™1000/2000 P2 | Illumina | San Diego, CA | 20046811 |
| Colibri™ Library Quantification Kit | Invitrogen | Carlsbad, CA | A38524500 |
| Quant-it dsDNA High-Sensitivity Assay Kit (Qubit) | Invitrogen | Carlsbad, CA | Q33120 |
| Sera-Mag SpeedBead Magnetic Carboxylate | Cytiva | Marlborough, MA | 65152105050250 and |
| Trypsin/Lys mix | Promega | Madison, WI | V5073 |
| TMTpro™ 16plex Label Reagent Set | Thermo Fisher Scientific | Waltham, MA | A44521 |
| C18 trapping column Reprosil-Pur | Dr. Maisch GmbH | Ammerbuch-Entringen, | r15.aq.0001 |

**Table. S3.**
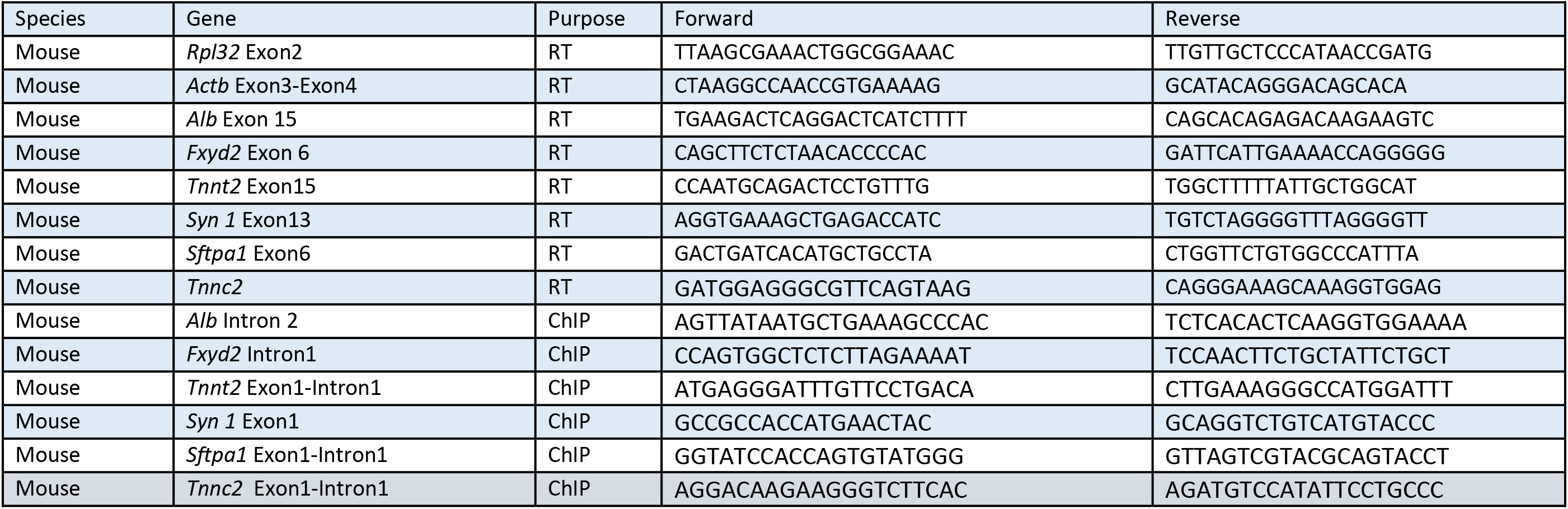
qPCR primers.

**Table. S4.** Antibodies.

| Antibody | Assay | Catalog No | Source | Manufacturer |
| --- | --- | --- | --- | --- |
| Pol II CTD (4H8) | ChIP-qPCR/seq | Sc-47701 | Mouse monoclonal | Santa Cruz |
| H3K27Ac | ChIP-qPCR/seq | 39133 | Rabbit polyclonal | Active Motif |
| H3 | ChIP-qPCR | PA-16183 | Rabbit polyclonal | ThermoFisher |
| H3K4m3 | ChIP-qPCR/seq | 39159 | Rabbit polyclonal | Active Motif |
| 5mC/m5C | MeDIP-seq/MeRIP-seq | AMM99021 | Mouse monoclonal 33D3 | Aviva System Biology |
| m6A | MeRIP-seq | 202-003 | Rabbit polyclonal | Synaptic System |
| IgG | ChIP-qPCR/seq | I-2000 | Mouse serum | Vector Laboratories |

**Table. S5.** PIXUL-Matrix protocols timeline.

| **Fresh frozen** | **ChIP-qPCR/seq** | **MeDIP-qPCR/seq** | **RT-qPCR/seq** | **MeRIP-qPCR/seq** |
| --- | --- | --- | --- | --- |
| CryoCore processing | 1hr |  |  |  |
| Isolation | Chromatin 2hrs | DNA 2hrs | RNA 4hrs | RNA 4hrs |
| Matrix | ChIP for qPCR 4hrs<br>ChIP for seq 16hrs | MeDIP-4hrs | RT 1.5 hrs | MeRIP 4hrs |
| Total time before PCR or library | 1 day- for qPCR<br>1.5 days for seq | 1day | 1day | 1.5 days |

| **FFPE** | **ChIP-qPCR/seq** | **MeDIP-qPCR/seq** | **RT-qPCR/seq** | **MeRIP-qPCR/seq** |
| --- | --- | --- | --- | --- |
| FFPE extraction | 2hrs |  |  |  |
| Isolation | Chromatin 1hr | DNA 2hrs | RNA 4hrs | RNA 4hrs |
| Matrix | ChIP for qPCR 4hrs<br>ChIP for seq 16hrs | MeDIP-4hrs | RT 1.5 hrs | MeRIP 4hrs |
| Total time before PCR or library | 1 day- for qPCR<br>1.5 days for seq | 1day | 1day | 1.5 days |

